# A quantitative intracellular peptide binding assay reveals recognition determinants and context dependence of short linear motifs

**DOI:** 10.1101/2024.10.30.621084

**Authors:** Mythili S. Subbanna, Matthew J. Winters, Mihkel Örd, Norman E. Davey, Peter M. Pryciak

## Abstract

Transient protein-protein interactions play key roles in controlling dynamic cellular responses. Many examples involve globular protein domains that bind to peptide sequences known as Short Linear Motifs (SLiMs), which are enriched in intrinsically disordered regions of proteins. Here we describe a novel functional assay for measuring SLiM binding, called Systematic Intracellular Motif Binding Analysis (SIMBA). In this method, binding of a foreign globular domain to its cognate SLiM peptide allows yeast cells to proliferate by blocking a growth arrest signal. A high-throughput application of the SIMBA method involving competitive growth and deep sequencing provides rapid quantification of the relative binding strength for thousands of SLiM sequence variants, and a comprehensive interrogation of SLiM sequence features that control their recognition and potency. We show that multiple distinct classes of SLiM-binding domains can be analyzed by this method, and that the relative binding strength of peptides in vivo correlates with their biochemical affinities measured in vitro. Deep mutational scanning provides high-resolution definitions of motif recognition determinants and reveals how sequence variations at non-core positions can modulate binding strength. Furthermore, mutational scanning of multiple parent peptides that bind human tankyrase ARC or YAP WW domains identifies distinct binding modes and uncovers context effects in which the preferred residues at one position depend on residues elsewhere. The findings establish SIMBA as a fast and incisive approach for interrogating SLiM recognition via massively parallel quantification of protein-peptide binding strength in vivo.

## INTRODUCTION

The proper function of cells depends on an enormous number of interactions between different proteins [1]. Interactions that are weak and transient are particularly important in controlling molecular events that are rapid and dynamic. Many of these interactions are mediated by peptide sequences known as Short Linear Motifs (SLiMs), which by definition do not form stable tertiary structures and instead are enriched in intrinsically disordered regions of proteins [2]. SLiMs bind to globular folded domains in their partners [2–5] and they can be recognized by a wide variety of modular protein domain families, with well-known examples including SH3, WW, and PDZ domains. Over 200 distinct families of globular SLiM-binding domains are known [2], and their binding to cognate SLiM peptides can control subcellular targeting, assembly of multi-protein complexes, and recognition of substrate proteins by modifying enzymes (e.g., kinases, phosphatases, ubiquitin ligases, etc.). SLiMs play key roles in signal transduction pathways and the control of protein stability [1, 5], and their gain or loss can drive the evolution of regulatory networks [6, 7]. Furthermore, SLiM-mediated interactions can be mimicked by viruses and other pathogens to co-opt host cell functions [8–11], they can be targets for drugs [12], and they may contribute to human diseases when dysregulated [13].

Major questions about SLiM function remain unresolved because of limitations in understanding how SLiM peptide sequences dictate their recognition (Fig 1A). “Consensus” residues, meaning those that are shared across most identified binding peptides, often play a central role in controlling the affinity of SLiMs for their relevant binding partners and specificity that minimizes off-target binding [3]. However, more-variable residues in adjacent non-core positions can also contribute substantially [14]. In addition, sequence features that affect the structural conformation of the peptide can modulate binding energetics without contributing to the domain-peptide interface [15]. Efforts to understand these binding determinants would benefit from improved methods for SLiM discovery and characterization. Current estimates suggest that roughly one third of the human proteome is intrinsically disordered [16, 17] and contains over 100,000 SLiMs [4, 18], with the majority yet to be identified. These SLiMs are estimated to fall into roughly 350 distinct classes, with over 4000 experimentally validated examples [2]. While some SLiM classes have been studied extensively, most have only a few known examples, and hence they lack accurate definitions of the range of functionally permissible sequences [2, 19]. Established consensus motifs usually lack information about which deviations from the consensus are functionally tolerated and they overlook contributions of flanking positions, which limits their utility for discovering novel motifs and for predicting the effects of polymorphisms or disease mutations. Furthermore, it is becoming increasingly evident that variation in the binding strength of SLiMs can tune the magnitude or timing of regulatory events, such as during cell cycle transitions and in the control of protein phosphorylation or degradation [20–24]. These findings highlight a critical need for comprehensive and quantitative approaches to define how variations in SLiM sequences affect their binding strength, specificity, and functional potency.

**Figure 1.**
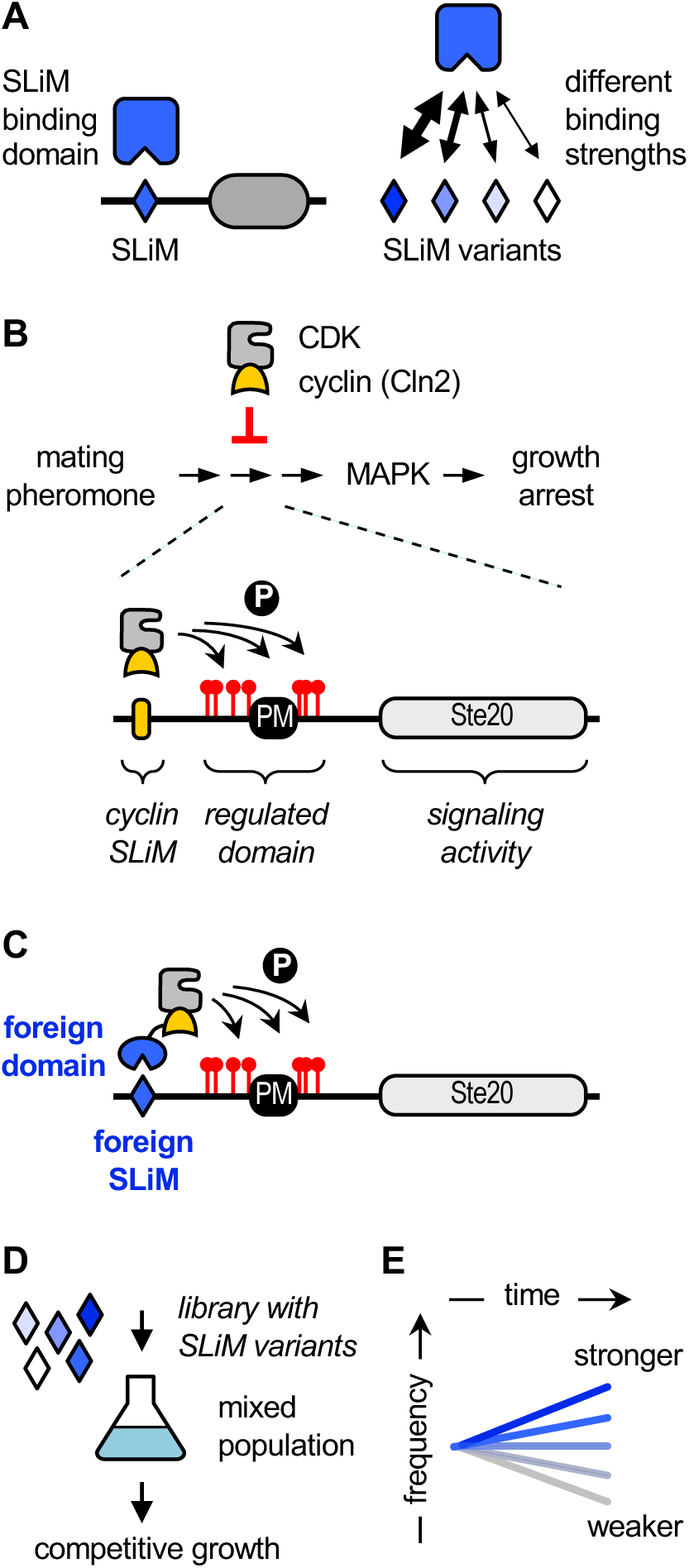
Overview of the SIMBA system. (A) Generic diagram emphasizing how SLiM sequence variants can differ in binding strength. (B) Top, CDK activity associated with the cyclin Cln2 blocks signaling through a pheromone-stimulated growth arrest pathway. Bottom, a chimeric signaling protein, Ste20^Ste5PM^, contains a plasma membrane binding domain (PM) and flanking phosphorylation sites from Ste5 joined to signaling domains from Ste20. A SLiM docking site for Cln2 promotes CDK phosphorylation of Ste20^Ste5PM^ at multiple sites, which inhibits signaling. (C) The role of cyclin docking can be replaced using foreign SLiMs and their binding domains, such that the foreign interaction allows cells to escape the growth arrest signal. (D) Libraries of Ste20^Ste5PM^ derivatives harboring large numbers of SLiM variants are introduced into yeast cells and analyzed by competitive growth. (E) The rate of SLiM enrichment or depletion is an indicator of binding strength.

A key attribute of SLiMs is that their short length (∼ 3-11 residues) and limited contact interface (e.g., only 3-4 residues buried in the binding pocket of their partner domains) means that their binding affinities are often relatively weak (e.g., K_D_ ∼ 1-500 μM) and rapidly dissociating [3, 25, 26]. Consequently, they mediate interactions that are inherently transient and dynamic. While this makes them well-suited for many important physiological functions (e.g., rapidly reversible interactions among signaling proteins), it can hinder their discovery by methods that rely on stronger binding, and it can complicate efforts to distinguish functional motifs from non-binding sequences. For example, tens of thousands of interactions in the human proteome have been identified via yeast two-hybrid screens, or by affinity purification followed by mass spectrometry [27–30], but in those screens SLiM-mediated interactions are statistically underrepresented, likely due to their weak affinities [5, 31]. Therefore, additional methods are needed to focus more specifically on interrogating SLiM-mediated interactions. Current approaches fall into two categories [32]: SLiM discovery (unbiased screens to identify new binding motifs), and SLiM characterization (analysis of sequence features that govern specific binding).

For the first category, SLiM discovery, high-throughput display-based technologies (phage display, bacterial display, and mRNA display) allow screening of peptide libraries that can exceed 10^8^ distinct sequences [33–40]. Despite providing a wealth of information on sequences that can bind a given bait domain, these approaches do not always yield robust information about the relative binding strength of the captured sequences or about features that prohibit binding [33]. The SLiM sequence features required for binding can be inferred indirectly from the most common residues in these sequences; while useful, these inferences are not comprehensive, as they provide no information about whether any residues absent from the identified sequences are compatible with binding, which is critical for full understanding of the binding determinants and for the accuracy of predictive modeling [41]. Importantly, to obtain affinity information about candidate SLiMs identified via screening, it is typical to conduct follow-up tests using low-throughput biophysical assays [19, 34, 42], which can introduce a bottleneck in post-screening stages. Therefore, alternate high-throughput methods to validate captured sequences and quantify their binding strengths would be valuable.

The second category, SLiM characterization, requires the initial identification of one or more binding peptides, which then can be probed for the sequence features that control their binding. These binding determinants can be defined comprehensively by using saturation mutagenesis, in which each position in a peptide is systematically varied to all possible amino acids. One common procedure for interrogating SLiM sequences uses “SPOT” arrays, in which peptides are synthesized in situ on a solid support [43, 44]. This method has the advantage of being accessible to many experimental labs as well as the option to incorporate residue modifications (e.g., phosphorylation) at specific peptide positions. However, the binding measurements are often only semi-quantitative, and they involve washing steps that disrupt equilibrium and bias results toward the strongest binders [3, 45]. A recent method, “MRBLE-pep”, uses peptide variants attached to spectrally coded beads to measure binding to purified domains in vitro [45]. This method can accurately quantify binding affinities for hundreds of peptides in parallel, although it requires specialized equipment and reagents as well as purified protein domains; moreover, the costs and effort scale in direct proportion to the numbers of domains to be tested, which can become prohibitive for larger-scale efforts in which the number of domains and peptide contexts range from dozens to thousands. Several additional approaches using functional assays in living cells have provided detailed characterization of peptides that act as kinase docking motifs [20, 46], transactivation domains [47] and degrons [48], although none has yet been generalized to permit their application to other SLiM-binding domain families.

All currently available methods for SLiM analysis have their own advantages and disadvantages [32]. Ideal approaches would integrate several key features: (i) a quantitative readout that directly correlates with biophysical properties (e.g., binding affinity) or functional characteristics (e.g., stability); (ii) scalability in terms of the number of domains and peptides that can be tested; and (iii) generalizability to accommodate a diverse range of domain and peptide classes. In addition, assaying interactions within a cellular environment can be desirable to replicate the conditions and effects of crowding found in the cytoplasmic milieu [49]. Therefore, to provide a new approach that can complement existing methods, help circumvent bottlenecks, and fill key gaps in knowledge, we have devised a high-throughput method that allows for systematic and quantitative analyses of SLiM binding using an in vivo assay. The approach is called “SIMBA”, for Systematic Intracellular Motif-Binding Analysis. By combining deep mutational scanning (DMS) with a competitive growth assay in which SLiM binding confers a growth advantage, we can quantify the relative binding strength of thousands of motif variants simultaneously and thereby define the rules of SLiM recognition with high accuracy. The work described here validates SIMBA methodology and initiates several downstream applications to demonstrate its feasibility and utility. We apply the approach to refine our understanding of three well-studied motif families by providing unique insights into the positive and negative contributions of non-core residues to binding strength, the impact of motif sequence context on residue preferences, and the interdependence of preferences at distinct motif positions. The findings highlight the potential for the SIMBA approach to substantially illuminate our current understanding of protein-protein interactions by providing a new analytical tool that can define recognition rules for large numbers of SLiMs and their binding domains.

## RESULTS

### Origin and Development of SIMBA Methodology

In general overview, the SIMBA method is an intracellular assay in which binding between a protein domain and a SLiM peptide regulates signaling through a yeast growth arrest pathway. In this system, cyclin dependent kinase (CDK) can block the growth arrest response by phosphorylating a protein in the MAPK-dependent signaling pathway (Fig 1B). To do so, the CDK requires one of its cyclins to recognize the target protein via a SLiM “docking site” (Fig 1B), which promotes dynamic, multi-site phosphorylation of the target [50, 51]. Because this phosphorylation inhibits the signaling protein [52], it allows cells to grow in the presence of the arrest signal (yeast mating pheromone). Previously, we exploited this antagonism of growth arrest to define the sequence requirements of docking peptides for binding to the native yeast cyclin, Cln2 [20]. Here, we have adapted this system to monitor binding between foreign globular domains and their SLiMs. To achieve this, we fuse the foreign SLiM-binding domain to yeast Cln2, and then use its cognate foreign SLiM peptide to replace the Cln2 docking motif in the CDK substrate (Fig 1C). As a result, binding of the foreign domain to its SLiM can drive substrate phosphorylation and block the growth arrest signal in a manner that reflects their interaction strength.

In the basic procedure, yeast cells contain two constructs. One encodes the foreign domain fused to Cln2, expressed from a galactose-inducible promoter. The other encodes the signaling protein that hosts the SLiM sequence to be tested; this protein is a chimeric signaling molecule, Ste20^Ste5PM^, that can be inhibited by CDK phosphorylation and can tolerate the insertion of peptide sequences [20, 50]. To monitor the effects of SLiM binding, expression of the domain-Cln2 fusion is induced, and then pheromone is added to activate the growth arrest pathway. There are two options for a quantitative readout. The first uses low-throughput assays of signaling, involving either transcriptional reporters or western blots that detect phosphorylation of a MAPK in the arrest signaling pathway, to quickly test small numbers of domain-peptide pairs. The second uses high-throughput assays, involving competitive growth of mixed cell populations, to screen libraries containing thousands of different SLiM sequences (Fig 1D). Here, stronger SLiM binding confers faster growth [20], and deep sequencing is used to analyze the rates of enrichment or depletion for all SLiMs in the population (Fig 1E), which are then converted to scores of relative binding strength (see Methods). It is worth emphasizing that, while our findings below and in subsequent sections indicate that in vivo strength correlates with binding affinity, the in vivo scores cannot be converted directly to a biophysical parameter (such as K_D_).

### Modular SIMBA components permit the study of diverse domains and peptides

A broadly applicable assay for peptide binding must have sufficient modularity to accommodate a diverse range of domain-peptide pairs. Initial tests using low-throughput assays showed that several distinct classes of SLiM-binding domain could be analyzed by the SIMBA method. We tested 6 SLiM-binding domains of 4 distinct structural types (Fig 2A, top), each of which was fused to the N-terminus of the yeast cyclin Cln2: an SH3 domain from the yeast protein Abp1 [53]; an Ankyrin Repeat Cluster (ARC) domain from human Tankyrase 2 (TNKS2^ARC4^) [54]; two WW domains from human YAP1 and NEDD4 [55, 56]; and two SWIB domains from human MDM2 and MDM4 [57, 58]. These domains were chosen because each had multiple known binding peptides with a range of affinities. Binding of these domains to their cognate SLiMs was readily detected by virtue of enabling the hybrid cyclin-CDK complex to block yeast pheromone signaling, as measured by MAPK phosphorylation or induction of a transcriptional reporter. For example, tests using the TNKS2^ARC4^ domain illustrated several key features (Figs 2B, S1A-C). First, the TNKS2^ARC4^ domain did not perturb the ability of Cln2 to recognize its own docking site (“LP_Ste5”), which provides a useful control for functionality of the fusion protein as well as an internal standard for binding strength. Second, the TNKS2^ARC4^ domain imparted the ability to inhibit signaling proteins containing its cognate SLiMs, which were not inhibited by Cln2 alone. Third, the magnitude of inhibition conferred by each SLiM peptide in vivo reflected their binding affinities measured previously in vitro. Fourth, the foreign SLiMs did not affect the levels of the recipient substrate protein but they did alter its electrophoretic mobility in cells expressing the TNKS2^ARC4^-Cln2 fusion (Fig 2B), consistent with inhibitory phosphorylation by the hybrid cyclin-CDK complex. Analogous results were obtained with the 5 other SLiM binding domains (Fig S1D-F), which are summarized in Fig 2A (bottom). Altogether, these results validate the utility of SIMBA as a modular and generalizable approach for detecting foreign domain-SLiM interactions and assessing their relative binding strengths.

**Figure 2.**
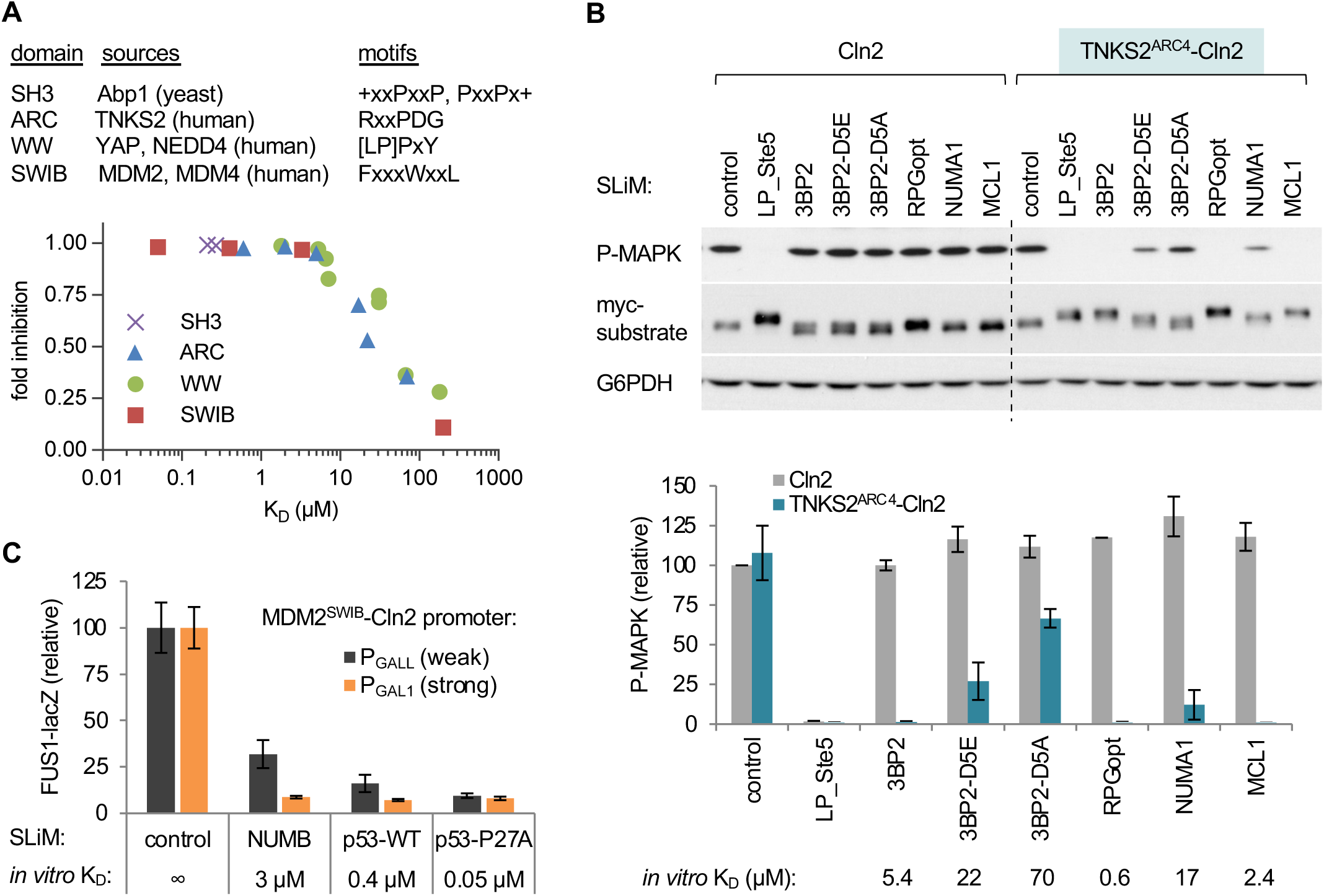
Test cases and low-throughput assays of SLiM binding via the SIMBA system. (A) Top, tested domains and their motif types. Bottom, summary of results presented in Figures 2B and S1D-F. The plot shows functional strength measured by SIMBA in vivo vs. previously measured affinities in vitro. Error bars are omitted for clarity. See also Figure S1C. (B) Cln2 and a TNKS2^ARC4^-Cln2 fusion were coexpressed with Ste20^Ste5PM^ derivatives harboring known TNKS2^ARC4^-binding peptides, and then pheromone signaling was assayed. Top, representive blots; SLiM binding blocks activation of the MAPK Fus3 (P-MAPK) and promotes a mobility shift of the myc-tagged Ste20^Ste5PM^ substrate. Bottom, quantification of results (mean ± SD; n = 2 [Cln2] or n = 6 [TNKS2^ARC4^-Cln2]), compared with in vitro binding affinity [54]. See also Figure S1A-B. (C) Improved resolution of strong interactions by expressing the MDM2^SWIB^ domain from a weaker promoter (P_GALL_) vs. the full strength promoter (P_GAL1_). Bars, mean ± SD (n = 4). See also Figure S1C.

The tests above showed that the SIMBA system can distinguish interactions with *K*_D_’s in the 5-100 μM range but starts to saturate for stronger affinities. To increase resolving power for these stronger interactions, we reduced the levels of the domain-cyclin fusion protein by using a weaker promoter to drive its expression. Specifically, we replaced the strong galactose-inducible promoter (P_GAL1_) with weakened versions, P_GALL_ and P_GALS_ (Fig S1G). Indeed, interaction strengths of peptides that bind the MDM2^SWIB^ domain with affinities in the 0.05-5 µM range were resolved better when the domain-Cln2 fusion was expressed from the weaker P_GALL_ promoter rather than the full-strength P_GAL1_ promoter (Fig 2C). Thus, simple adjustments of the system’s sensitivity can permit resolution of affinities spanning roughly three orders of magnitude (*K*_D_ = 0.05-100 μM). In the following sections, we will present the results from high-throughput experiments that validate the utility of the SIMBA approach for screening large numbers of domain-peptide pairs as well as for revealing unforeseen determinants of binding strength and specificity.

### SIMBA allows comprehensive screening of binding strength and preferences in vivo

To confirm that the functional strength in vivo reflects biochemical binding affinity, we established a test case involving the TNKS2^ARC4^ domain. We chose this domain because its binding affinity had been measured in vitro for almost 200 SLiM peptides, including a set of all 152 single-site substitutions in an 8-residue peptide (RSPPDGQS) derived from human 3BP2 [54]. Thus, we used SIMBA to measure binding strengths for this same set of single-site variants (Fig 3A), and then compared the results from our in vivo assay to those from the prior in vitro measurements. This comparison revealed a good agreement for the full set of 153 variants (i.e., WT plus 152 mutants) (Figs 3B-C). Moreover, the SIMBA results recapitulated the distinct categories of selectivity at individual peptide positions that were previously seen in vitro (Figs 3C, S2A); examples include an exclusive requirement for Arg at p1, broad tolerance with a continuum of binding strengths at p2, and unique intolerance for a Pro residue at p7. We also mutagenized two extra residues on either side of the 8-residue motif (i.e., p-2, p-1, p9, and p10) and confirmed that these flanking regions have minimal influence on binding (Figs 3A-B).

**Figure 3.**
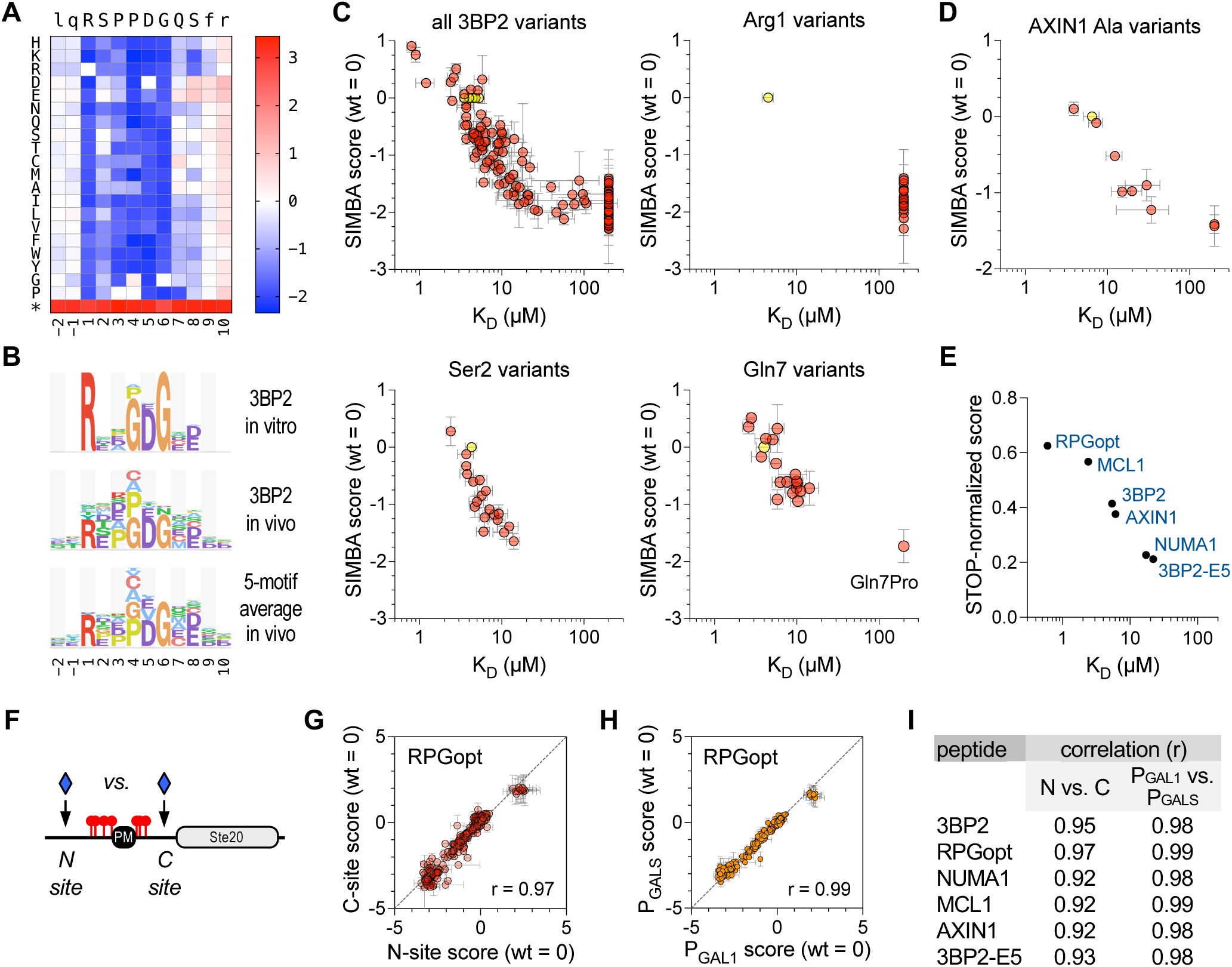
Interrogation of TNKS2^ARC4^-binding peptides by SIMBA. (A) Heatmap showing effects of all single-position substitutions in the 3BP2 motif. Colors denote enrichment (red) or depletion (blue) relative to the wild-type motif, calculated as log scores by Enrich2 software (see Methods). The asterisk denotes a termination codon. The data average four independent experiments using P_GAL1_-driven TNKS2^ARC4^-Cln2 (two each of N-site and C-site libraries). (B) Logos showing TNKS2^ARC4^ sequence preferences observed for the 3BP2 motif in vitro [54] and in vivo (this study), plus the average in vivo results from 5 distinct motifs (see also Figure S2B-D). For clarity, only positive preferences are shown (see Methods). (C) Plots of SIMBA scores vs. K_D_ for all 3BP2 variants as well as representative individual positions. Yellow, wild-type; pink, missense variants. SIMBA scores are mean ± SEM (n = 4). K_D_ values [54] are the mean of technical duplicates (n = 1 ± SE of the regression fit); K_D_ values that were not quantifiable in vitro were assigned values of 200 µM to allow inclusion in the plot. See also Figure S2A. (D) SIMBA scores vs. K_D_ values [54] for the wild-type AXIN1 peptide plus 9 Ala mutants, plotted as in panel (C). See also Figure S2B-D. (E) Comparison of STOP-normalized SIMBA scores to in vitro K_D_ [54] for 6 wild-type peptides. See also Figure S2B-D. (F) Diagram of N-site and C-site locations for inserting SLiM sequences. (G) Correlation of SIMBA scores for variants of the RPGopt motif in N-site vs. C-site locations. Data are the mean ± range (n = 2). See also Figure S2E. (H) Correlation of SIMBA scores for RPGopt motif variants when the TNKS2^ARC4^-Cln2 fusion was expressed from different strength promoters (P_GAL1_ vs. P_GALS_). Data are the mean ± SEM (n = 4; two each of N-site and C-site libraries). See also Figure S2F. (I) Summary of Pearson correlation coefficients (r) for the indicated pairwise comparisons. See also Figures S2E-F.

Because all peptide-encoding plasmids are pooled together and assayed simultaneously during the competitive growth, it is minimal extra work to increase the number of peptides tested by several fold. Therefore, we performed DMS on five other parent peptides concurrently with the 3BP2 peptide (Fig S2B-C), each of which had been assayed individually in vitro [54]. These peptides included 3 native sequences from other proteins (NUMA1, MCL1, AXIN1), a previously defined variant with optimum residues at each position (RPGopt), and a weakened version of the 3BP2 peptide with a Glu substitution at p5 (3BP2-E5). In total, binding was measured in parallel for 1374 total peptides. The sequence preferences averaged across 5 motifs (excluding the 3BP2-E5 mutant) resembled that obtained from the 3BP2 motif alone (Fig 3B), although we observed context-specific features that will be described below. None of the peptides showed strong preferences at positions flanking the core 8-residue motif (Fig S2C). For the AXIN1 peptide, binding affinities were measured previously for Ala mutations at 9 positions (p-1 to p8) [54]; as we found with the 3BP2 variants, these AXIN1 mutations affected binding strength similarly in vivo and in vitro (Fig 3D). The score distributions from the 6 parent peptides reflected their expected binding strengths. Namely, those with stronger affinities (Fig S2B) gave broader variant score distributions and larger (in absolute value) negative scores (Fig S2D), indicating that the wild-type peptide is further from non-functional. Also, as described previously [20], stronger peptides more closely approach the maximum possible score achieved by mutants containing termination codons (Fig S2D, red), which have the greatest selective advantage because they eliminate the protein that mediates the growth arrest signal. When the parent peptide scores were normalized to these maxima, their in vivo strengths correlated with their previously measured affinities (r = 0.93) (Fig 3E). Thus, the good agreement between in vivo and in vitro binding strengths is observed when testing either mutant variants of individual peptides or groups of distinct parent peptides.

Finally, we asked if the SIMBA results might be influenced by the regional polypeptide context in which the SLiM peptide was embedded, by comparing results of inserting them at two distinct locations in the recipient protein – either on the N-terminal or C-terminal side of the phosphorylated region (Fig 3F). The results in the two location contexts were strongly correlated (Figs 3G, 3I, S2E). Therefore, the peptide motifs are functionally autonomous, with binding specificities that are independent of the surrounding polypeptide context. We also compared results when expressing the TNKS2^ARC4^ fusion protein from the full-strength (P_GAL1_) versus a weakened (P_GALS_) promoter (Fig S2A), and we found robust agreement (Figs 3H-I, S2F-G). Even with the full-strength promoter, strong peptides that were indistinguishable in low-throughput transcription or MAPK phosphorylation assays, such as 3BP2 (5 µM) and RPGopt (0.6 µM) (see Figs 2B, S1A), showed resolved binding strengths in the competitive growth assay (Figs 3E, S2D), and both reductions and increases in binding strength were detectable for mutant variants of each peptide (Fig S2C-D). Thus, sub-micromolar affinities of peptides do not preclude discovery of their binding determinants. We speculate that the resolution of strong binders is improved in the growth assay because the longer timespan of the experiment allows small differences to be reinforced and compounded. Collectively, our findings show that SIMBA can serve as an accurate gauge of relative biochemical affinity, that it reveals the same motif sequence preferences as would be observed in vitro with purified components, and that these preferences are independent of the host protein context of the SLiM or the expression level of its partner domain.

### Local context effects revealed by comparing multiple parent peptides

As mentioned earlier, we performed DMS for multiple TNKS2^ARC4^-binding peptides. When their residue preferences were grouped by position, it became evident that some parent peptides had strikingly different preferences at p4 (Fig 4A-B). Namely, for three peptides (3BP2, RPGopt, and NUMA1), Pro and Gly were equally the most favored residues at p4 (Fig 4C), in agreement with the previous in vitro results using the 3BP2 peptide [54]. In contrast, for the peptides from MCL1 and AXIN1, Gly was among the most disfavored residues at p4 (Fig 4C). These two peptides also showed a unique preference for Pro at p2 (Fig 4A), and both are inherently Pro-rich (Fig 4B), suggesting that they might be predisposed to form a type II polyproline helix (PPII helix). This left-handed structure possesses a restricted conformational flexibility that can reduce the entropic cost of binding [59, 60], and hence we hypothesize that this energetic benefit is disrupted when a Gly residue is introduced. In support of this interpretation, the main-chain trajectories for all peptides co-crystallized with TNKS2^ARC4^ show nearly identical left-handed topology from p1 through p5 [54], compatible with a PPII helix (Fig 4D-E).

**Figure 4.**
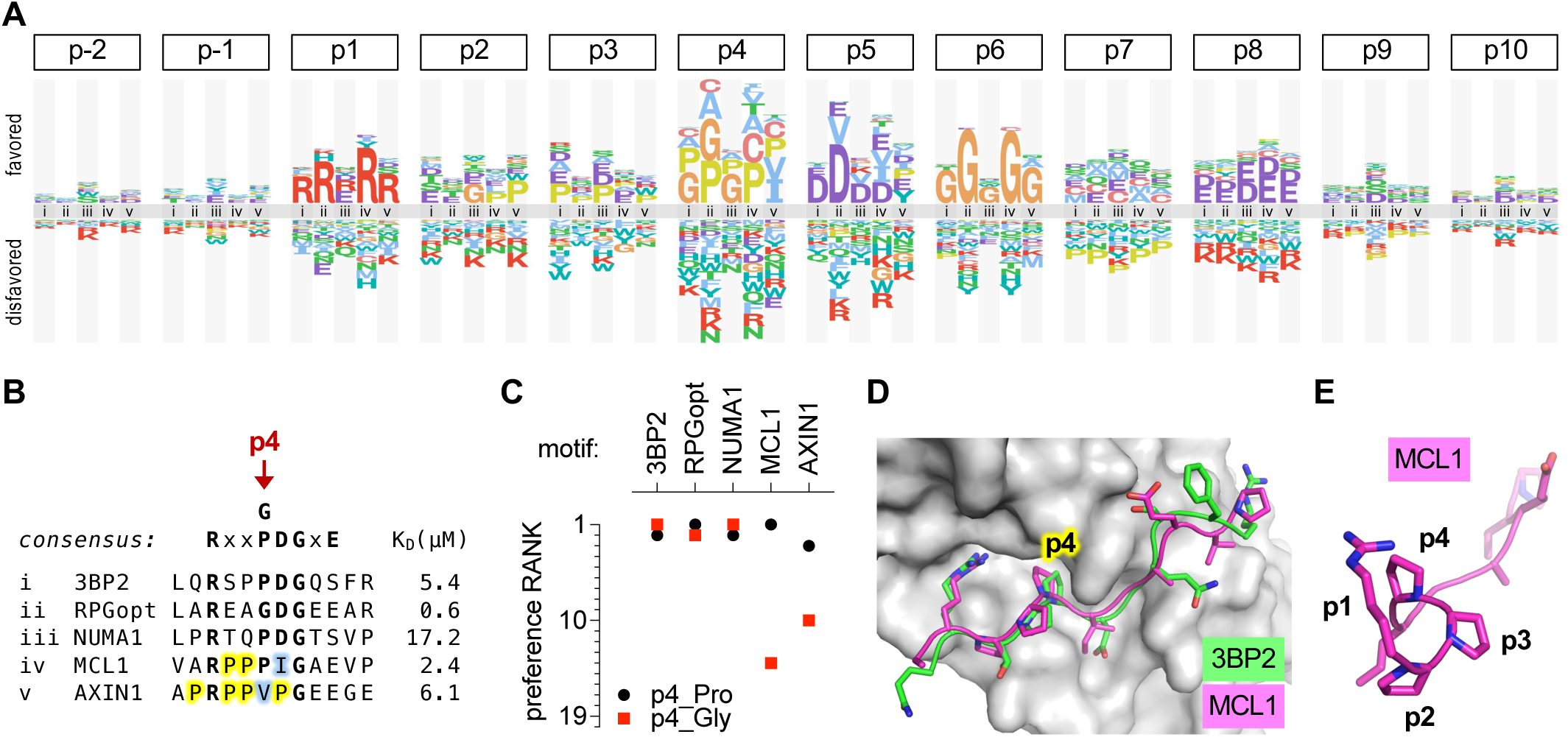
Local context affects residue preferences at some positions. (A) Logo comparing the residue preferences at each peptide position in the context of 5 different parent peptides that bind TNKS2^ARC4^ (labeled i-v; see panel [B]). (B) Sequence and K_D_ of parent peptides (i-v) analyzed in panel (A). Yellow, Pro residues flanking p4 in MCL1 and AXIN1. Blue, residues that are atypically preferred in MCL1 and AXIN1. (C) Preference ranks of Pro and Gly at p4 in 5 different parent motifs. (D) Similar trajectories and p4 contacts of 3BP2 and MCL1 peptides bound to TNKS2^ARC4^ [54]. PDB IDs: 3twr, 3twu. (E) Rotated view of the TNKS2^ARC4^-bound MCL1 peptide, showing left-handed trajectory of poly-proline sequence from p2 to p4

Another notable context effect at p4 was the unusual preference for Val or Ile in the AXIN1 peptide (Fig 4A). Because Val at p4 had been observed to be disfavored in the 3BP2 context [54], it was assumed that the same would be true in the AXIN1 context; hence, to explain strong binding by the AXIN1 peptide, it was hypothesized that the negative impact of Val at p4 was offset by the presence of a favored Glu residue at p8 [54]. Surprisingly, however, our experiments revealed that Val at p4 is not suboptimal in the AXIN1 context and instead it is one of the two most favored residues (Fig 4A). Conceivably, because p4 in AXIN1 is flanked on both sides by Pro residues, Val or Ile might maintain the predisposition toward PPII structure while improving packing against the hydrophobic p4-binding pocket in TNKS2^ARC4^. Similarly, the native residues at p5 for MCL1 (Ile) and AXIN1 (Pro) are not as disfavored in each of these parent peptides as they are in the 3BP2 peptide context, and instead they are favored as strongly as the more-common Asp residue (Fig 4A). In contrast to these context-dependent differences, it is noteworthy that the requirement for Gly at p6 and the severe intolerance for Pro at p7 were observed in all parent peptides (Fig S2C). Collectively, our findings illustrate the informative value of performing DMS analysis on multiple parent peptides, as it can reveal context-dependent preferences that would not be suspected from analysis of any single peptide motif. Such benefits emerge readily from the SIMBA methodology due to its ability to analyze thousands of peptide sequences simultaneously.

### Defining SLiM recognition rules for the SWIB domain from human MDM2

As an additional test case, we used SIMBA to characterize the sequence preferences of the MDM2^SWIB^ domain, which drives ubiquitin-mediated degradation of the tumor suppressor p53 and is a target of anticancer compounds designed to block its peptide-binding pocket [61, 62]. We performed DMS on two human MDM2^SWIB^ binding peptides, one from p53 and another from NUMB, that have a shared core motif (FxxxWxxL) but different affinities (Fig S1E). The SIMBA results revealed similar preference patterns for both peptides (Fig 5A), including strong selectivity for hydrophobic residues at the three core positions p5, p9, and p12 (FxxxWxx[LIVMF]) that engage a deep hydrophobic cleft in the MDM2^SWIB^ domain [57, 63]. Additional preferences were evident at the non-core positions, most obviously for bulky aromatic or nonpolar residues at p8. There was a clear difference in the optimization of the two peptides, as binding to the NUMB peptide could be strengthened by numerous mutations (Fig 5B), particularly by replacing suboptimal residues with preferred residues at several non-core positions (p4, p8, p13, p15) (Fig 5A). In contrast, the p53 peptide contains preferred residues at p4 and p8, providing a potential explanation for why, despite identical core motifs, the binding affinity of the p53 peptide is roughly ten-fold stronger than that of the NUMB peptide (Fig S1E) [33]. The data also revealed strong negative preferences that likely relate to peptide conformation rather than binding contacts between the peptide and domain. Namely, in both peptides, Pro and Gly were largely disfavored from p7 to p13, consistent with their propensity to disrupt the α-helical conformation of the bound peptide [57, 63]. Interestingly, p6 tolerates Pro, despite being within the α-helical region. At p13, which immediately follows the core motif, Pro was the most disfavored residue in both peptides (Fig 5A). This is notable given that Pro is the wild-type residue at this position in the p53 peptide, and its replacement with Ala strengthens p53 binding affinity in vitro [15, 64, 65]. In the NUMB peptide, a Pro mutation at p13 results in a severe loss of MDM2^SWIB^ binding, emphasizing that the context of the starting peptide influences its robustness to mutation.

**Figure 5.**
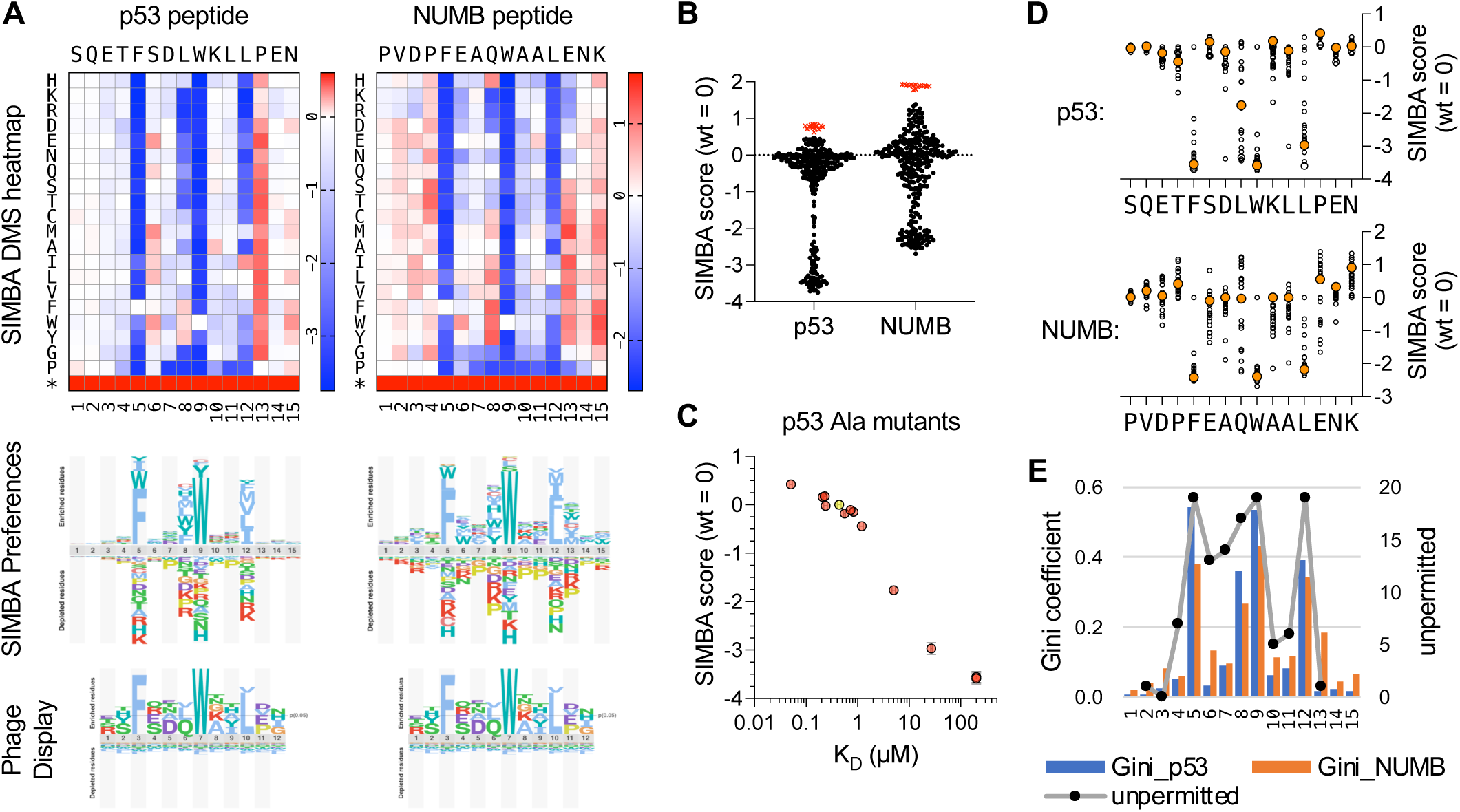
Determinants of binding strength for MDM2^SWIB^ peptide ligands. (A) Top, heatmaps of DMS results for p53 and NUMB peptides (mean, n = 2, for each peptide). Middle, sequence preference logos derived from the SIMBA scores. Bottom, logo from the ProP-PD database (https://slim.icr.ac.uk/proppd/) [33] showing residue frequencies in peptides captured by MDM2^SWIB^ phage display; this logo is shown twice to facilitate comparison with the SIMBA preferences in each of the two peptide contexts above. (B) Distribution of SIMBA scores for all missense variants (black circles) and the average STOP codon (red X symbols) at the 15 positions in each of the 2 parent motifs. (C) SIMBA scores vs. K_D_ values [65] for the wild-type p53 peptide (yellow) and 12 Ala mutants (pink). K_D_ values that were not quantifiable in vitro were assigned as 200 µM to allow inclusion in the plot. (D) Plot of SIMBA scores for Ala residues (large, orange-filled circles) vs. all other residues (small, unfilled circles) at each position in the p53 and NUMB peptides. (E) Bars show Gini coefficients, calculated from normalized SIMBA scores, for each position in the p53 and NUMB peptides. Black dots connected by a grey line show previous scoring of the number of unpermitted substitutions at each position of an optimized MDM2^SWIB^-binding peptide (MPR<u>F</u>MDY<u>W</u>EG<u>L</u>N) [66].

The effect on MDM2^SWIB^ binding affinity has been measured for alanine substitutions at 12 positions along the p53 peptide [65]. A comparison of these affinities with their corresponding SIMBA scores showed excellent agreement (r = 0.91) over a range of 3-4 orders of magnitude in K_D_ (Fig 5C), giving further confirmation that SIMBA provides an accurate measure of relative binding strength. Compared to alanine scanning, DMS data provide a more nuanced definition of how binding is dictated by sequence. In particular, we observed that non-core positions can substantially modulate binding strength (Fig 5D), but this effect can be overlooked by Ala-scanning for two reasons: (i) the effect of Ala is often more neutral compared to other residues; and (ii) the effect of Ala depends on the favorability of the wild-type residue being replaced (e.g., Ala at p8 is more disruptive in p53 than in NUMB because the wild-type residue is more favorable in p53 [Leu] than in NUMB [Gln]). The range of residue tolerance at each position, as revealed by these DMS measurements, can be summarized using the Gini coefficient as a measure of inequality (Fig 5E). It shows a pattern that is largely similar for both the p53 and NUMB peptides and is also consistent with a pattern of “not permitted” substitutions (defined as > 3-fold change in IC_50_) observed in previous mutagenic scanning of an optimized MDM2-binding peptide [66]. Finally, a comparison to previous phage display data for the MDM2^SWIB^ domain [33] (Fig 5A, bottom) similarly illustrates how SIMBA results can provide more refined discrimination between the degrees of preference at distinct positions, as well as information about substitutions that result in the loss of binding (which is absent in phage display results). Altogether, these analyses of MDM2^SWIB^ binding preferences reinforce the utility of the SIMBA method for providing accurate, high-resolution definitions of motif binding determinants and for understanding the sequence basis for differences in peptide binding strength.

### Specificity determinants and context effects in WW domain-binding peptides

To further explore how SIMBA methodology could be used to interrogate SLiM preferences and context dependence, we studied two different WW domains that bind peptides with a common core motif, [LP]PxY, or “PY peptides”. We chose the first WW domain from human YAP (YAP^WW1^) and the third WW domain from human NEDD4 (NEDD4^WW3^), to allow comparison with prior information about their peptide ligands, binding affinities, and structural details of peptide-domain contacts [55, 56, 67–73]. Notably, statistical analyses of peptides captured by these two domains in phage-display experiments suggested there were correlations between residue preferences at different positions in the peptide [74]. That is, the most favored residue at a given peptide position might depend on the identity of residues at other positions. We sought to conduct direct empirical tests of such contingent preferences by performing DMS on multiple parental peptides that differ in their starting sequence context.

Initial tests using low-throughput SIMBA assays and a small number of peptides (Fig S1F) confirmed that WW domain binding was readily detected for higher-affinity peptides (K_D_ ∼ 2-7 µM) and was weak or undetectable for lower-affinity peptides (K_D_ ∼ 50-180 µM). For subsequent high-throughput assays and DMS, we chose 22 parental peptides from three categories (Fig S3A): (i) 15 natural peptides reported to bind one or both WW domains; (ii) 3 mutationally-optimized peptides (UGR2, UGR1, UGR1-LYG) that bind the NEDD4^WW3^ domain with strong, sub-micromolar affinity; and (iii) 4 peptides representing the consensus sequences of possible motif sub-classes (PYcon1-4) that were suggested by the statistical analyses noted above [74]. We used the competitive growth assay to test binding of these 22 parental peptides to each WW domain (Fig 6A-B). To compare their binding strengths against each other (rather than comparing mutant variants with a single wild-type sequence), we calculated z-scores that express the magnitude of enrichment of all peptides relative to a set of “nonbinder” control sequences (see Methods). The two domain fusions showed similar scores for the control Cln2-binding motifs (LP peptides) (Fig 6A), which provide internal standards for their binding capacities. Peptides derived from viral capsid proteins were weak binders (Fig 6B), consistent with prior in vitro data (Fig S3A). Many of the other peptides were recognized by both WW domains, but the rank order of binding strength was clearly distinct for the two domains, and some peptides showed striking specificity for one domain over the other (Figs 6A-B). Of note, those with the strongest specificity for YAP^WW1^ have a Pro residue immediately following the core Tyr residue (Fig 6B). From here forward, we will refer to this core Tyr position as p0 and denote other positions by their distance before (p-2, p-1, etc.) or after (p+1, p+2, etc.).

**Figure 6.**
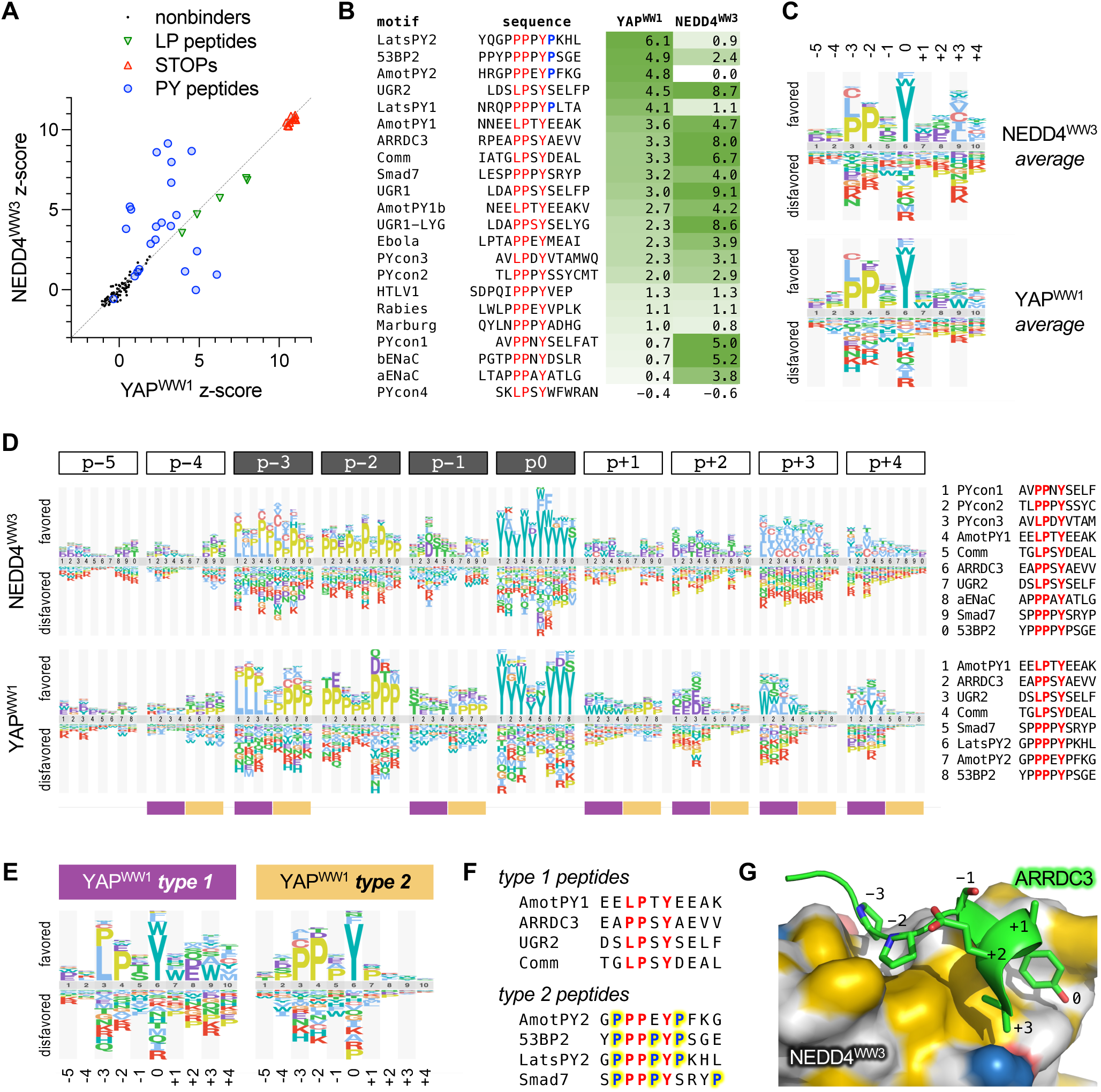
Specificity determinants and context effects in WW domain-binding peptides. (A) Scatterplot of z-scores (median, n = 6) for peptides tested for YAP^WW1^ and NEDD4^WW3^ binding. (B) Sequences and binding z-scores (median, n = 6) for 22 peptides, sorted by YAP^WW1^ binding strength. The core [LP]PxY motif is colored red, and Pro residues at p+1 are colored blue. (C) Logos showing average sequence preferences for each WW domain. (D) Logos comparing preferences at each peptide position in the context of multiple parent peptides (identified at right for each WW domain). At bottom, plum and tan bars mark the distinct type 1 and 2 preference patterns summarized in panel (E). (E) Logos showing distinct YAP^WW1^ preferences for type 1 and 2 peptides. (F) Sequences of the type 1 and 2 peptides that contribute to the logos in panel (E). Yellow highlighting and blue font mark Pro residues at non-core positions in type 2 peptides. (G) AARDC3 peptide bound to the NEDD4^WW3^ domain (PDB ID: 4n7h) [72]. Hydrophobic regions on the NEDD4^WW3^ surface are colored yellow [105].

For DMS, we mutagenized twelve of these parental peptides at 12 positions each, including the core motif plus flanking residues on both sides (Fig S3B). Collectively, these experiments assayed binding of each WW domain to 2748 peptide sequences. The averaged sequence preferences from all parent peptides (Fig 6C) matched the expected pattern at the core positions (i.e., [LP]PxY), while also suggesting milder preferences at surrounding positions. However, these averages obscured distinct context-dependent patterns that became evident when the preferences were grouped by position (Fig 6D). Especially striking was a bifurcation of YAP^WW1^-binding peptides into two classes, which we designated as types 1 and 2 (Fig 6E), that show distinct requirements at p-3: type 1 tolerates either Leu or Pro at p-3, whereas type 2 strongly favors Pro over Leu. Type 2 motifs also showed a unique preference for Pro at non-core positions p-4, p-1 and p+1 (Fig 6D-E). Because the parental type 2 peptides are especially Pro-rich (Fig 6F), their partiality toward Pro suggests a hypothesis similar to the one raised earlier for TNKS2^ARC4^ peptides. Namely, the type 2 sequences are likely predisposed to form a PPII conformation that is adopted by peptides in the bound state [68, 71–73, 75], which reduces the entropic cost of binding, and hence substitutions that disrupt this PPII propensity are disfavored.

On the C-terminal side of the core motif, type 1 peptides preferred acidic residues at p+2 as well as nonpolar residues at p+3 and p+4 (Fig 6D). These preferences fit with prior structural studies that identified contacts between C-terminal non-polar residues and a hydrophobic pocket on the NEDD4^WW3^ domain (near the Tyr-binding pocket) [68, 71, 72, 76]. In such cases, a right-handed helical turn immediately following the p0 Tyr allows a nonpolar side chain at p+3 or p+4 to occupy the hydrophobic pocket (Figs 6G, S3C), and acidic side chains at p+2 can stabilize the helix by making intrapeptide hydrogen bonds back toward p-1. Similar C-terminal preferences were strikingly absent in type 2 peptides (Fig 6D-E), perhaps because the Pro at p+1 might prevent the helical turn and/or occupy the hydrophobic pocket itself, as seen in the complex of YAP^WW1^ with the Lats_PY2 peptide [73] (Fig S3C). Indeed, for YAP^WW1^, Pro was the most favored p+1 residue in type 2 peptides but the least favored in type 1 peptides (Fig 6D). For type 1 peptides, YAP^WW1^ and NEDD4^WW3^ showed subtle differences in their favored nonpolar residues at p+3 (Fig 6D). Specifically, YAP^WW1^ showed a notable tolerance for the aromatic residue Trp whereas NEDD4^WW3^ generally preferred aliphatic residues (I/L/V/A/C), perhaps reflecting differing composition of their hydrophobic pockets. The inclusion of Cys with these nonpolar residues is consistent with prior findings that it interacts favorably with hydrophobic membrane or protein environments [77–79], and that it ranks as nonpolar in several hydrophobicity scales [80].

### Contingent preferences implicate contributions from intrapeptide interactions

We also noticed that preferences at p-1 were influenced by peptide context (Fig 6D). For example, the most favored p-1 residues in several peptides were Ser or Thr while in other peptides they were Asp or Ala. Furthermore, in the Smad7 context there was an unusual preference for aliphatic residues (I/L/V). We reasoned that these preferences might be influenced by the C-terminal flanking sequence (C-flank), because peptides that adopt a C-terminal helical turn can form intrapeptide interactions in which the p+2 side chain projects back toward p-1 (Figs 6G, S3C) [68, 70, 72], and the Smad7 peptide adopts an atypical hairpin-like conformation that places the Pro residue at p+4 near p-1 (Fig S3C) [67]. To address this possibility, we appended seven different C-flank sequences to the core motifs from three parental type 1 peptides, and then tested the p-1 preferences for all 21 combinations (Fig 7A). Although a few of these hybrid peptides bound YAP^WW1^ too weakly to be informative, the majority indicated that the C-flanks had clear effects on p-1 preferences of both WW domains. Namely, with some exceptions, the general trends were that two of the C-flanks (PYcon3, aENaC) imposed a p-1 preference for Asp, four others (Comm, AmotPY1, ARRDC3, UGR2) imposed a preference for Ser or Thr, and one (Smad7) imposed a distinct preference for Pro. Notably, the C-flanks that imposed the Asp preference at p-1 have a Thr residue at p+2, whereas those that imposed the Ser/Thr preference have a Glu residue at p+2 (Fig 7A). In light of the aforementioned intrapeptide interactions between p+2 and p-1 seen in some structures, we tested if changing the residue identity at either position altered preferences at the other (Fig 7B). Although the effects were not as strongly determinative as when entire C-flanks were swapped, we found that the identity of the p+2 residue did alter the rank of p-1 preferences in several motif contexts (Fig 7B, left). Namely, for three type 1 motifs (UGR2, aENaC, PYcon3), the presence of Thr rather than Glu at p+2 led to an increased preference at p-1 for Asp rather than Ser/Thr. In contrast, in the type 2 motif from Smad7, the p+2 residue had little effect on its unusual p-1 preferences. In reciprocal tests (Fig 7B, right), the identity of the p-1 residue influenced p+2 preferences modestly for two motifs (ARRDC3, Comm), in which the presence of Asp rather than Ser at p-1 led to an increased preference for Thr at p+2. Such influences were more variable for the aENaC motif and absent for the strong UGR2 motif. Collectively, our observations reveal pair-wise contingent preferences between two positions (p-1 and p+2) that are further influenced by the surrounding context. Because p-1 and p+2 residues remain solvent accessible in type 1 peptides (Fig S3D), rather than buried as part of the peptide-domain interface, their coordinated effects on binding likely signify a role in intrapeptide interactions that stabilize the bound conformation.

**Figure 7.**
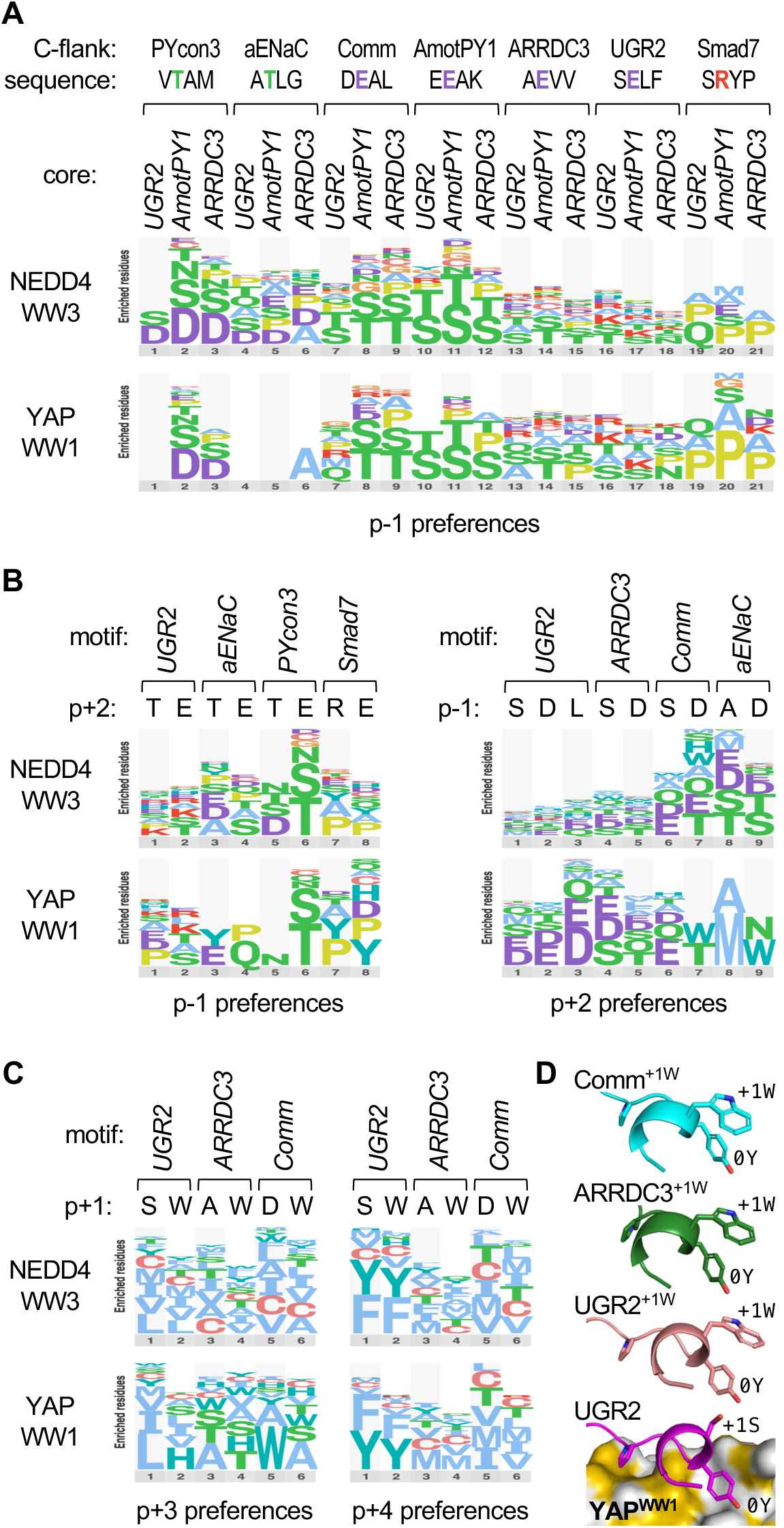
Contingency relationships between distinct peptide positions. (A) Logos of p-1 preferences in 21 different peptide contexts. Seven 4-residue C-flank sequences were appended to each of three core motifs (…[LP]PxY), and then the p-1 codons were randomized and tested for binding YAP^WW1^ and NEDD4^WW3^. For clarity, the logos show only the most preferred residues, and they omit data from peptides for which the difference in z-score between the tested WW-Cln2 fusion and unfused Cln2 was less than 1. (B) Logos showing preferences at p-1 when the p+2 residue is varied (left), or preferences at p+2 when the p-1 residue is varied (right). (C) Logos showing preferences at p+3 or p+4 when the p+1 residue is varied. Data in panels A-C are derived from the median synonym z-scores from 3 independent experiments. (D) Predicted conformations of peptides bound to YAP^WW1^. Bottom, prediction for the UGR2 peptide, showing 3 side chains for reference (-2P, 0Y, +1S). Above, analogous predictions for peptides with p+1 Trp substitutions (+1W), aligned with the bottom structure and hiding the YAP^WW1^ domain for clarity. The Trp groups are predicted to lie next to the p0 Tyr groups (0Y) with their planar faces in a perpendicular geometry.

Curiously, type 1 peptides showed a substantial preference for Trp at p+1 (Fig 7D-E), a position that normally faces away from the WW domain in peptides with the C-terminal helical turn (Fig 6G). We considered the possibility that a Trp at p+1 provides an alternative way to favorably occupy the hydrophobic pocket, which could thereby preclude the role for nonpolar residues in the C-terminal helix. However, replacing p+1 with Trp did not alter the preferences for nonpolar residues at p+3 or p+4 displayed by NEDD4^WW3^, or those at p+4 displayed by YAP^WW1^, although it did noticeably shift the p+3 preferences shown by YAP^WW1^ (Fig 7C). These results suggest that the contributions to binding strength conferred by Trp at p+1 and nonpolar residues at p+3/p+4 are additive, not competitive. This finding favors an alternative explanation in which the p+1 Trp helps stabilize the type 1 C-terminal helix, perhaps via an aromatic-aromatic interaction [81] with the adjacent p0 Tyr. Indeed, structural predictions using AlphaFold [82] suggest that these Trp side chains lie next to the Tyr in a perpendicular geometry (Fig 7D) that is common for aromatic-aromatic interactions [81]. Altogether, our interrogations of context-dependent preferences reveal that different peptide conformations and domain pocket characteristics can contribute to a remarkable variety of subtly distinct binding modes that are not adequately described by average preferences or those of any singular motif.

### Preferences at non-core positions influence predictions of binding strength

Consensus sequences describe the minimal required features of binding motifs, but due to their relatively low complexity there can be multitudes of matching sequences that are not easily distinguishable. For the consensus sequence [LP]PxY, there are 1730 matches in the intrinsically disordered regions of the human proteome. To rank their potential for binding the YAP^WW1^ or NEDD4^WW3^ domains, we used the comprehensive SIMBA data to derive position-specific scoring matrices (PSSMs) that quantitatively weight the preference for every possible residue at each position in the extended motif. Then, these PSSM values were summed over the length of the motif for each of the 1730 human sequences. This process was performed separately for each WW domain and using PSSMs derived from all motifs combined versus from only type 1 or only type 2 motifs. Each PSSM dispersed the 1730 matches into a broad distribution (Fig 8, “hits”), which is primarily due to variation at non-core positions, as the core consensus describes only 2 distinct sequences (i.e., LPxY and PPxY). For comparison, we also calculated PSSM sums for the parental motifs that had been tested in SIMBA experiments (Fig 8, “tested”). The strong binders among these tested motifs were generally in the top quartile of PSSM sum distributions. Interestingly, for YAP^WW1^ the type 1 PSSM gave very high scores for the strong type 1 motifs but low scores for the strong type 2 motifs, whereas the type 2 PSSM gave the reverse pattern. These type-specific results predict that only sequences yielding PSSM sums within the top 10% of the distribution would show binding strengths comparable to those tested in our experiments. For NEDD4^WW3^, the different PSSMs all gave high scores to the strongest binders, yet we again observed that some of the type 1 motifs were underestimated by the type 2 PSSM and vice-versa (Fig 8, “tested”). Overall, these results illustrate that quantitative PSSMs derived from SIMBA data can help distinguish strong binders from bulk consensus matches, and they also emphasize the importance of using PSSMs specific for each distinct binding mode to achieve the best assessment of candidate sequences. By extension, without prior knowledge of distinct binding modes, evidence for their existence could emerge from disagreements between observed binding strengths and predictions that are based on a single PSSM.

**Figure 8.**
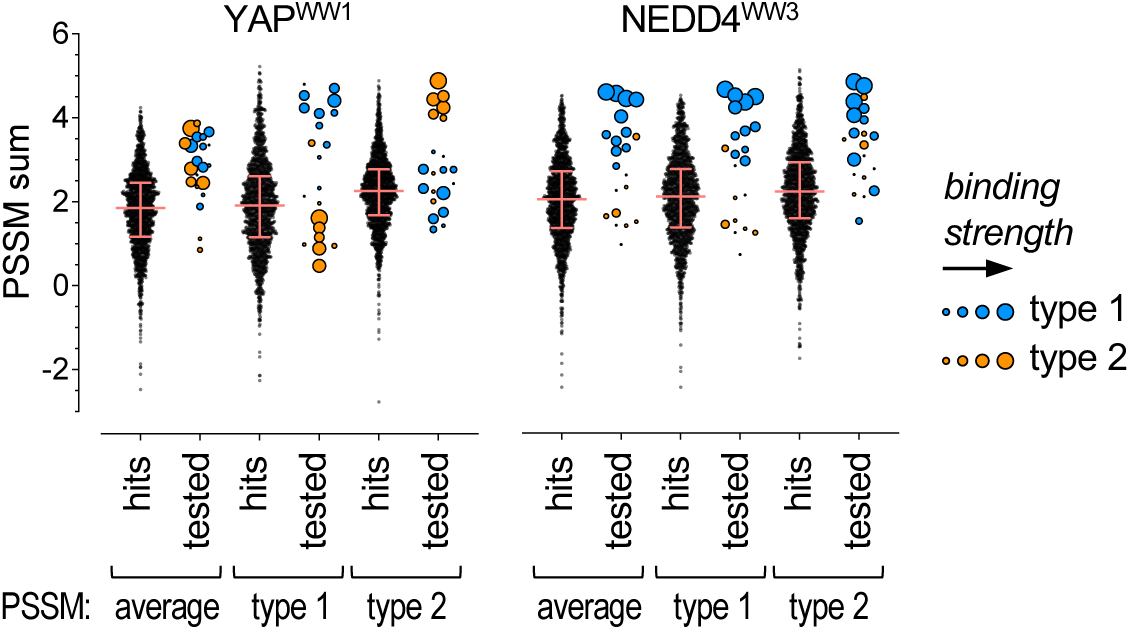
Stratification of [LP]PxY matches by PSSM scoring. All 1730 matches to the [LP]PxY consensus sequence (“hits”) in disordered regions of the human proteome were scored using 6 different PSSMs. For each of 2 WW domains, 3 PSSMs were derived from the DMS results: one from all motifs (average), one from type 1 motifs only, and one from type 2 motifs only. The PSSM values for individual residues were summed across 10 motif positions (xx[LP]PxYxxxx) to calculate a predicted score (PSSM sum) for all sequences. For all hits, the distribution of sums obtained using each PSSM is shown; pink lines denote median and quartile values. For all motifs tested by SIMBA (“tested”), PSSM sums were calculated similarly, and their symbol sizes are proportional to their observed SIMBA scores (see Figure 6B).

## DISCUSSION

In this study we have described and validated a new method, SIMBA, for performing systematic and quantitative analyses of SLiM-mediated binding to multiple distinct domains. This comprehensive and quantitative analysis provides a fast, easy, and low-cost alternative to traditional biochemical assays of protein-peptide binding. The power of the approach lies in its ability to rapidly quantify the binding strength of thousands of variant motifs in parallel and thereby reveal the specificity determinants for the queried domain. Further advantages include that it monitors binding in an in vivo setting, it can discern variations in binding strength over a broad range, and it is amenable to both low-throughput and high-throughput assays.

Previous studies on docking peptides for cyclins and MAPKs provide precedents that competitive growth assays in yeast can be used for fine-scale measurement of peptide binding strength [20, 46]. SIMBA now generalizes these strategies in a way that can be applied to a broad range of different types of domains and peptides, without depending on their normal biological functions. We envision that this method will be of primary utility in two categories of investigation. The first is to define the sequence determinants of SLiM binding strength and specificity for a given domain or group of related domains. The second is to compare the relative binding strengths of large sets of candidate SLiM peptides. Such candidates might emerge from other screens (e.g., phage display hits) or they might be potential binding peptides identified in the proteome as matches to an existing “consensus” sequence for a specific domain. In either case, SIMBA allows dozens to thousands of such candidates to be analyzed quickly and easily.

SIMBA provides a valuable complement, though not a substitute, to other methods for determining biochemical affinity. Its measurements of binding strengths are relative rather than absolute. For many purposes, such as for defining the sequence preferences of SLiM-binding domains, this relative binding information of mutant variants is sufficient, and it is not necessary to know the affinity of the wild-type motif. In cases where there are pre-existing measurements of K_D_ values for some peptides that bind a given domain, their inclusion in SIMBA experiments allows them to serve as benchmarks against which to compare the many other peptides being tested.

The findings in test cases described here clearly demonstrate how systematic analyses of SLiM sequence requirements can illuminate unanticipated complexities in binding determinants even for well-studied examples such as TNKS2^ARC4^, YAP^WW1^, and NEDD4^WW3^. Although we had primarily expected to confirm previously defined binding rules for these domains, in each case we also obtained evidence for different binding modes and contingent relationships between different motif positions that could not have been predicted based on prior knowledge. These observations illustrate the intrinsic discovery potential of the SIMBA approach that emerges readily from its ease of performing massively parallel analyses. Namely, gathering empirical binding results for large numbers of domain-peptide pairs can provide unexpected insights that would be missed in more limited analyses (e.g., those focusing on singular peptide contexts).

The SIMBA method is fundamentally a variation of the classic yeast two-hybrid system [83, 84], except the interaction being tested controls a phosphorylation reaction rather than transcription. In principle, either method could be used to investigate SLiM binding determinants, and the most crucial advances for large-scale interrogation arise from the combination of competitive growth and deep sequencing to allow massively parallel analyses. Nevertheless, SIMBA offers several additional benefits that are notable. (i) It can detect weak binding in the 1-100 μM range that includes many physiologically relevant SLiM interactions [3]. (ii) The interaction controls multisite substrate phosphorylation that yields a continuum of regulatory strength [52], which makes it sensitive to binding strength over a broad range. (iii) The phosphorylation reaction in vivo is dynamic and rapidly reversible [85], which can help ensure that the measured output reflects binding at equilibrium. (iv) Because the fused cyclin (Cln2) can still recognize its own docking site, it provides an internal standard against which to compare the strength of binding mediated by the fused domain. (v) The effects of the SLiM interaction are under acute control, as they require induced expression of the cyclin fusion and an external stimulus, and hence there is no chronic selection pressure to skew population distributions before the experiment begins. (vi) It allows adjustments in the required binding strength (by using different strength promoters) so that multiple affinity ranges can be monitored.

Because SIMBA monitors enrichment or depletion of all sequences in the tested population, the quality of data is similar for both functional and nonfunctional motifs. This aspect is a notable contrast to capture-based approaches, such as phage display, that provide only indirect inferences about any sequences that were not captured. Dependable quantification of a broad range of binding strengths, from strong binders to non-binding sequences, as well as identification of prohibited residue substitutions, will be of great value in developing accurate ranking and filtering criteria for future efforts at in silico prediction of binding motifs. High quality, comprehensive binding data should also be useful for structural modeling and for predicting pathogenic impacts of sequence variants. For example, quantitative data from large numbers of peptides, including abundant examples of non-binders, can provide training information to improve computational predictions of protein complexes and their binding affinities [86–88]. Separately, they can provide empirical tests of binding effects of cancer-associated mutations in the COSMIC database [89] and improve the confidence in pathogenicity predictions generated by AlphaMissense [90], especially for variants in disordered regions of proteins that lack the structural features used as a main input of that algorithm.

Finally, in addition to providing mechanistic insights into binding specificity, SIMBA could also accelerate development of useful tools for basic biochemical research or synthetic biology. For example, the ability to rapidly identify variant peptides with distinct binding strengths could allow the design of degron tags, enzyme docking sites, or localization anchors whose effects on recipient target proteins can be finely tuned over a broad range. Separately, SIMBA approaches could also assist with identifying competitive peptide inhibitors of SLiM-binding domains. Such peptide inhibitors could serve as drug proxies or guides for peptidomimetic compounds [91] and could also be used to pre-screen the effects of target inhibition in human cell lines [92]. In applications currently underway, we have already applied the SIMBA method to over thirty other SLiM-binding domains, and we have extended the approach to include unbiased screens for discovery of new binding motifs. Thus, we expect that SIMBA, due to its ability to systematically define SLiM recognition rules and functional potency en masse in vivo, will be a versatile tool for numerous areas of future investigation.

## EXPERIMENTAL PROCEDURES

### Yeast Strains and Growth Conditions

Standard procedures were used for growth and genetic manipulations of yeast [93, 94]. Unless indicated otherwise, cells were grown at 30°C in yeast extract/peptone medium with 2% glucose (YPD), or in synthetic complete medium (SC) lacking histidine and/or uracil with 2% glucose or raffinose. Yeast strains and plasmids are listed in Tables S1 and S2, respectively. To construct plasmids encoding fusions of Cln2 to foreign SLiM-binding domains, the domain sequences were amplified by PCR or obtained as synthesized gene fragments, and then they were inserted as BamHI-XmaI fragments between the GST and CLN2 sequences of a P_GAL1_-GST-CLN2 construct [50]. All such constructs used for competitive growth assays harbored a full-length CLN2 fragment (residues 1-545), whereas many initial constructs used for low-throughput signaling assays contained a truncated fragment (residues 1-372). Related constructs harbored weakened promoters [95] called P_GALS_ (-415 to -5) and P_GALL_ (-431 to -5), which were amplified by PCR and inserted in place of the full-strength promoter P_GAL1_ (-675 to -5). We note that the expression levels were reproducibly greater with P_GALS_ than with P_GALL_, which is the reverse of the rank strength reported previously [96]. Strains used for competitive growth assays were constructed by integrating P_GAL_-GST-domain-CLN2 constructs into the genome at the *HIS3* locus of strain PPY2617; PCR was used to confirm single copy integration. These strains were transformed with plasmids that encode derivatives of a chimeric signaling protein, Ste20^Ste5PM^, that harbor different SLiMs [20].

### Signaling Assays

For low-throughput assays of SLiM binding, we used methods similar to those described previously [20, 50] to monitor the ability of domain-Cln2 fusion constructs to inhibit pheromone signaling, either by using a transcriptional reporter (*FUS1-lacZ*) or by immunoblotting for phosphorylated MAPK (Fus3). Plasmids encoding Ste20^Ste5PM^ chimeras with various SLiMs were either (i) co-transformed with a domain-Cln2 fusion plasmid into a *STE5-8A ste20Δ* strain (PPY2617); or (ii) transformed into a similar strain containing an integrated copy of a relevant domain-Cln2 fusion plasmid. Measurement of *FUS1-lacZ* transcriptional induction and MAPK activation followed methods described previously [20, 97]. Cell cultures were grown to exponential phase in SC liquid medium containing 2% raffinose and lacking histidine and/or uracil. Cyclin expression was induced by adding 2% galactose for 90 minutes. For *FUS1-lacZ* assays, cells were treated with pheromone (50 nM) for 45 min, and β-galactosidase activity was measured as described previously [97]. To assay MAPK activation, cells were treated with pheromone (50 nM) for 15 min, whole cell lysates were prepared as described previously [98], and then proteins were resolved by SDS–PAGE and transferred to PVDF membranes (Immobilon-P; Millipore). Primary antibodies used include mouse anti-phospho-p44/42 (1:1000, Cell Signaling Technology #9101), rabbit anti-phospho-p44/42 (1:5000, Cell Signaling Technology #4370), rabbit anti-myc (1:200, Santa Cruz Biotechnologies #sc-789), and rabbit anti-G6PDH (1:100000, Sigma #A9521). Horseradish peroxidase-conjugated secondary antibody was goat anti-rabbit (1:3000, Jackson ImmunoResearch #111-035-144). Enhanced chemilluminescent detection used a BioRad Clarity substrate (#170-5060). Densitometry was performed using ImageJ software. Quantified measurements of signaling outputs using either assay are plotted relative to samples in which a control peptide (SNGNGSGSNGN) [20] is present in the Ste20^Ste5PM^ chimera. Fold inhibition is the difference in signal between the test peptide and the control peptide, as a fraction of the control signal.

### Library Construction and Competitive Growth Assays

The construction of mutant libraries and the performance of bulk growth competition assays were based on procedures described previously [20]. SLiM variants were tested in the context of a chimeric Ste20^Ste5PM^ signaling protein, in which the membrane-binding domain from Ste5 and its flanking CDK phosphorylation sites replace the native membrane-binding domain in Ste20, and SLiMs are placed at either the N or C terminal side of the Ste5 fragment [20, 50].

To construct libraries of SLiM variants, SLiM-encoding sequences were amplified by PCR using oligonucleotides as templates and primers that anneal to flanking linker sequences (Gly Gly Ser Gly). The template oligonucleotides were designed such that single codons were randomized (NNN) to generate all 64 nucleotide variants. All templates for a given motif were pooled in equal amounts and then amplified by 10 or 12 PCR cycles (98° for 10 sec; 56° for 20 sec; 72° for 6 sec) using different primer sets for insertion at the N-terminus (ggagtgacgtcGGAGGTAGTGGT and ctcacgctagcTCCAGATCCACC) or the C-terminus (actatacgcgtGGAGGTAGTGGT and tctttgcatgcTCCAGATCCACC). Column-purified PCR products were digested with restriction enzymes (AatII and NheI for insertion at N terminus; MluI and SphI for insertion at C terminus), treated with calf intestinal phosphatase, and then ligated (16 hrs at 18°, then 10 min at 65°) into the appropriately digested vector (pPP4375 or pPP4745). The ligation products were transformed into E. coli (XL-10 Gold Ultracompetent Cells; Agilent/Agilent Technologies) and plated on LB+Amp plates. Colonies (exceeding the number of variant sequences per library by > 10-fold) were harvested by adding LB+Amp liquid and gently agitating with a glass rod spreader, and then plasmid DNA was prepared from the suspension of pooled colonies. The isolated DNAs were checked by Sanger sequencing to verify that all four nucleotides were comparably represented at each position in the randomized codon. To construct plasmids with individual peptide motifs (e.g. wild type), the oligonucleotides used as PCR templates were unique sequences without codon randomization, or for some early constructions the insert fragments were obtained by annealing complementary oligonucleotides as described previously [20].

For competitive growth assays, a solution with equimolar amounts of each individual codon library was prepared. A mixture of control plasmids containing wild-type SLiMs plus unrelated and random sequences was spiked into the solution (making up 1-2.5% of the total) to create the final pool, which was transformed into yeast strains containing the relevant P_GAL_-domain-CLN2 fusions. After the transformation procedure, the bulk of the transformation mixture was split and plated onto two -URA plates, plus a diluted aliquot (1%) was plated onto a third -URA plate. Colonies on the diluted plate were counted to ensure that the number of transformants exceeded the number of mutant variants in the library by at least 10-fold. Colonies on the concentrated plates were suspended in 10 mL of –URA/raffinose liquid medium, washed twice with 20 mL -URA/raffinose, then diluted into 50 mL -URA/raffinose to yield a density of ∼ 4 × 10^6^ cells/ml (OD_660_ ∼ 0.3). These cultures were incubated in a shaking water bath for 4 hrs, then diluted back to OD_660_ ∼ 0.01 and incubated overnight for ∼16 hrs. The cultures were diluted back again (to OD_660_ ∼ 0.6 in 50 mL), incubated for 1.5-2.5 hrs, and treated with 2% galactose for 75 min (in a volume of 70 mL) to induce cyclin expression. At this time (t = 0), an aliquot (∼ 38 mL, 3 × 10^8^ cells) was harvested, and the remaining culture was treated with pheromone (500 nM) and returned to incubation. Cells were diluted with fresh medium (including galactose and pheromone) after the first 8 hr and subsequently after every 12 hr to maintain an OD_660_ below 1 (in a volume of 50 mL). Additional aliquots (∼ 20 mL, 3 × 10^8^ cells) were harvested at 8, 20, 32 and 44 hr. Harvested cells were collected by centrifugation (5 min., 3200 rpm, room temp), washed with 10 mL sterile water, resuspended in 1 mL sterile water, and transferred to 1.5 mL microcentrifuge tubes. These suspensions were centrifuged, the supernatants were aspirated, and the pellets were frozen using liquid nitrogen and stored at -80°C.

### DNA Preparation and Deep Sequencing

Plasmids were isolated from yeast cell pellets following previously described methods [20, 99]. DNA was purified using the Zymo Research ZR Plasmid Miniprep Kit (#D4015). Frozen cell pellets were thawed and suspended in 200 µl of solution P1 and treated with 7 μL Zymolyase (0.2 units/μL; Zymo Research #E1005) at 37°C for 1.5 hr, with vortexing approximately every 20 mins to disperse clumps, before proceeding with the remaining steps. Samples of plasmid DNA (4 μL) were subjected to PCR (17 cycles, 50 μL total volume) with primers that include standard P5 and P7 sequences for binding to Illumina flow cells during next generation sequencing (NGS). The forward primer included the P5 sequence, followed by an Illumina sequencing primer binding site, a 6-nucleotide bar code, and an upstream plasmid-annealing sequence; the reverse primer included a P7 sequence, followed by a 6-nucleotide i7-index sequence, an i7 sequencing primer binding site, and a downstream plasmid-annealing sequence. Aliquots (5 μL) of the PCR products were run in 1 % agarose gels to confirm the presence of the desired product. For any timepoint samples with low product yield, PCRs were repeated using NotI-digested plasmid and/or increased PCR cycles (19-25). Gel band intensities were quantified by densitometry (ImageJ), and then equal amounts of products from all timepoints for a given strain were pooled, purified using Zymo Spin I columns (Zymo Research #C1003-250), and eluted in 30 μL of 10 mM TRIS-HCl, pH 8. The concentration of the eluted products was measured using NanoDrop spectrophotometry, and then equal amounts of samples from all strains were pooled, gel-purified in triplicate from a 1% gel (NEB Monarch DNA Gel Extraction Kit #T1020L), eluted in 20 μL, and the triplicate eluates were combined. The final concentration was measured by Qubit fluorometry and the product size distribution was verified by Fragment Analyzer (Agilent) before being sent for Illumina-based NGS (paired-end sequencing, 150 bp) by a commercial vendor (Novogene).

### Sequencing Data Analysis

NGS data were de-multiplexed by strain and timepoint using bar code and index identifiers. To compare mutant variants versus wild-type SLiM sequences, we used Enrich2 software [100] to obtain read counts for all sequence variants. Then, we added 0.5 counts to any variants depleted to 0 counts by the 32-hr timepoint (to prevent Enrich2 from ignoring fully-depleted variants), and used Enrich2 to calculate fitness scores that describe the rate of change of mutant variants compared to the wild-type sequence [20, 100]. All scores reported here were calculated from 32-hr time courses.

To compare groups of sequences where there was no single wild-type reference standard, such as when comparing multiple WW domain binding peptides to each other, we calculated their enrichment relative to a set of non-binder control sequences. For this, we used 2FAST2Q [101], a program in Python, to count the occurrence of sequences from all strains and timepoints, converted each value to a frequency by dividing by the total counts, and normalized each frequency relative to the starting (t=0) frequency. We calculated a raw score for each timepoint as the log2 of the relative frequency divided by the time in hours (raw score = log2[relative frequency]/time). For subsequent calculations we used the median raw score from all timepoints for a given sequence. We defined a set of non-binder controls as all 236 missense variants of the core Tyr codon in four [LP]PxY motifs, plus 9 other non-[LP]PxY sequences. Then, we calculated z-scores as (X-µ)/σ, where X is the raw score of a test sequence, µ is the mean raw score of the non-binder set, and σ is the standard deviation of the non-binder raw scores. Hence, this z-score represents the number of standard deviations that a test sequence differs from the mean of non-binders.

To create logos of sequence preferences, we transformed Enrich2 scores or z-scores into a preference metric that expresses the bias for each residue at a given motif position relative to the set of all possible residues, following procedures described previously [20]. For DMS analyses of wild-type motifs, the raw Enrich2 score for each amino acid variant was normalized to the lowest (= 0) and highest (= 1) scores in a given motif array: (normalized score) = (raw score – minimum) ÷ (mean terminator score – minimum). Then, the normalized scores were converted to a frequency metric by dividing each by the sum of all scores at the same position: (frequency) = (normalized score) ÷ (column sum). Finally, the frequency metric was converted to a preference metric by subtracting 0.05, so that a neutral preference is represented by zero, favored residues are positive, and disfavored residues are negative: (preference score) = (frequency – 0.05). These preference scores were used to generate sequence logos via a web-based tool (http://slim.icr.ac.uk/visualisation/index.html). For experiments that probe how positional preferences in WW domain binding peptides are affected by altered peptide context (such as swapping C-terminal flanking sequences), the calculated z-scores for all variants in the control strain (unfused Cln2) were subtracted from the corresponding scores in the test strains (WW-Cln2 fusion). These Cln2-subtracted z-scores for all non-terminator variants were normalized separately for each position that was individually interrogated. Then, these normalized scores were converted to frequency metrics, preference scores, and sequence logos as described above. The plotted results exclude peptides for which the Cln2-subtracted z-score was less than 1. To generate a preference logo from in vitro affinities of 3BP2 variants, we first calculated the inverse of published K_D_ values [54], assigned an inverse value of zero to variants with unquantifiable K_D_’s, and then used these inverse values to calculate frequency and preference metrics as described above.

PSSMs for scoring matches to the [LP]PxY consensus sequence were derived as follows. First, as described above, normalized scores were calculated for each WW-binding motif. These values were then transformed to a difference PSSM [20] by subtracting the average of all residues at a given position from the value of each residue at that position: (difference score) = (residue score) – (position average). Then, the difference scores were averaged for subsets of motifs, as listed in Figure 6D: all motifs (sequences 0-9 for NEDD4^WW3^ or 1-8 for YAP^WW1^), type 1 motifs (sequences 1-8 for NEDD4^WW3^ or 1-4 for YAP^WW1^), and type 2 motifs (sequences 9 and 0 for NEDD4^WW3^ or 5-8 for YAP^WW1^). Finally, to give identical boundaries for all PSSMs, the averaged difference scores were further transformed such that positive values were normalized to the maximum (= 1) and negative values were normalized to the minimum (= -1). Human proteome sequences matching the [LP]PxY consensus were obtained using the SLiMSearch algorithm [102]. For each sequence, the corresponding boundary-normalized PSSM values for each residue at each position were summed across 10 motif positions (xx[LP]PxYxxxx) to obtain a predicted score (PSSM sum). PSSM sums were calculated for reference sequences in the same way.

For competitive growth assays of SLiM variants, fitness scores and standard errors were calculated by Enrich2 software [100]. Other statistical analyses, including calculation of means, medians, SD, SEM, Pearson’s correlation (r), and Gini index, were performed using Microsoft Excel. The numbers of biological replicates are described in the Figure Legends.

### Structure representations, analysis, and predictions

Illustrations based on prior crystallography or NMR structural data were generated using PyMOL software and original Protein Data Bank (PDB) coordinates. Calculations of buried surface area of peptide residues bound to WW domains were performed using PISA, an online protein interface analysis tool (https://www.ebi.ac.uk/msd-srv/prot_int/pistart.html). Structural predictions of peptide-domain complexes were generated by AlphaFold 2 [82], implemented using ColabFold [103] and UCSF Chimera X software [104]; illustrations were created in PyMOL.

## Data availability

All necessary data are available in the submitted manuscript or supporting materials. Large datasets derived from competitive growth assays and next generation sequencing, including variant counts and calculated enrichment scores, have been deposited at Mendeley Data: (https://doi.org/10.17632/nghf59hf4s.1).

## Supporting information

This article contains supporting information.

## Funding and additional information

This work was funded by grants from the NIH to P.M.P. (R01GM057769 and R01GM145795). The content is solely the responsibility of the authors and does not necessarily represent the official views of the National Institutes of Health. N.E.D. and M.O are funded by a Cancer Research UK Senior Cancer Research Fellowship (C68484/A28159) and by a UKRI grant EP/X042065/1 and Boehringer Ingelheim Fonds travel grant to M.O.

## Abbreviations

The abbreviations used are

SLiM: short linear motif
SIMBA: systematic intracellular motif binding analysis
DMS: deep mutational scanning
CDK: cyclin dependent kinase
MAPK: mitogen activated protein kinase
PSSM: position-specific scoring matrix
NGS: next generation sequencing
ARC: ankyrin repeat cluster
GST: glutathione S-transferase.

## SUPPORTING INFORMATION

### Supporting Figure Legends

**Figure S1.**
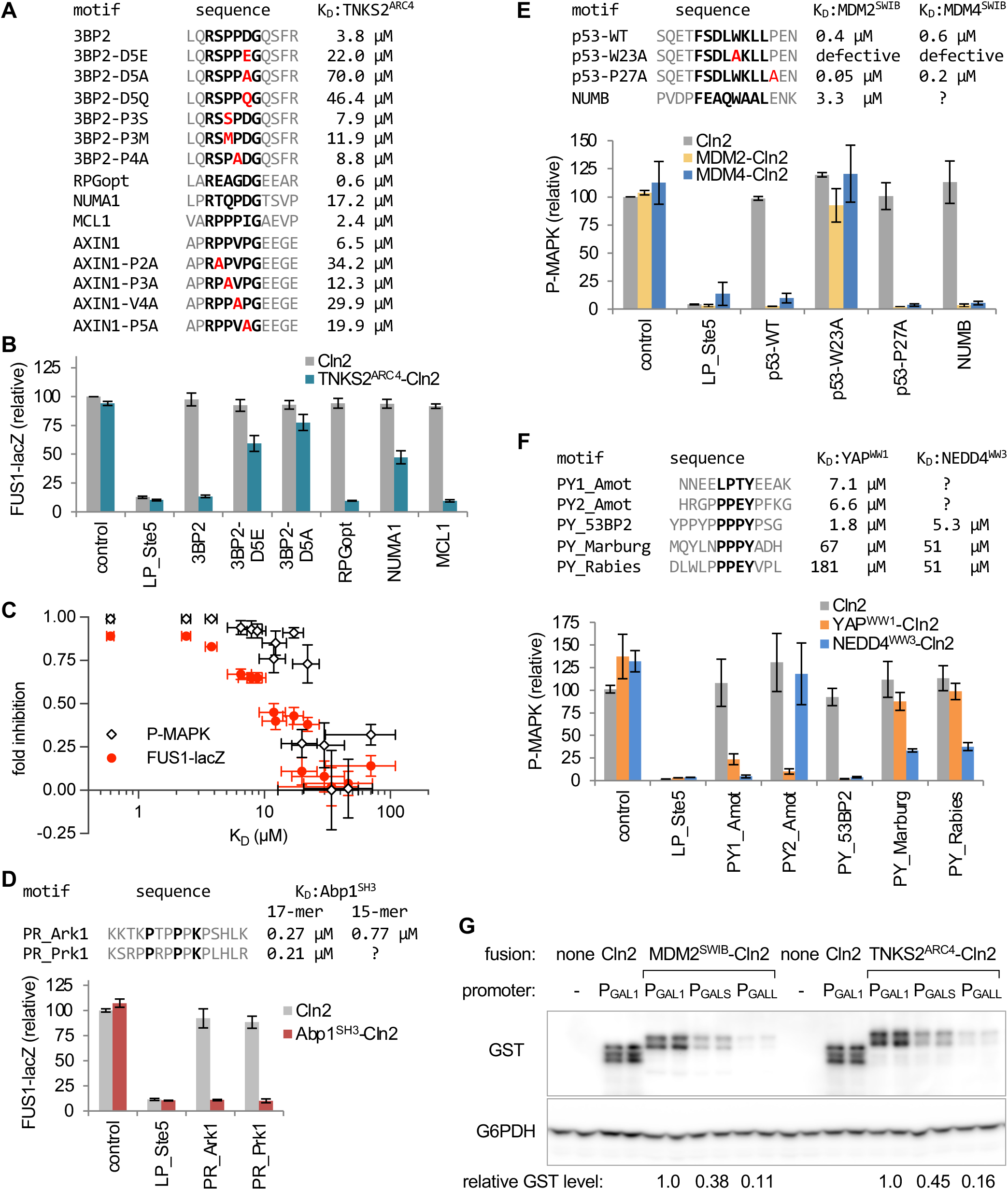
Low-throughput assays of SLiM binding to six domains. (A) List of tested TNKS2^ARC4^-binding peptides. K_D_ values are from [54]. (B) The experiment shown in Figure 2B was repeated using the transcriptional reporter, FUS1-lacZ (mean ± SD; n = 4). (C) Comparison of the fold inhibition in FUS1-lacZ vs. P-MAPK assays, for all peptides in panel A. Both assays show affinity-dependent inhibition, but the P-MAPK assay detects weak binding with greater sensitivity. Fold inhibition (mean ± SD, n = 4) is the difference in signal between the test peptide and the control, as a fraction of the control signal. K_D_ values are mean ± SE [54]. (D) SLiM recognition by the Abp1^SH3^ domain, assayed using the FUS1-lacZ reporter. Bars, mean ± SD (n = 3). K_D_ values from [53]. (E) SLiM recognition by MDM2^SWIB^ and MDM4^SWIB^ domains, assayed as in Figure 2B. Bars, mean ± range (n = 2). K_D_ values from [33, 65]. (F) SLiM recognition by YAP^WW1^ and NEDD4^WW3^ domains, assayed as in Figure 2B. Bars, mean ± range (n = 2). Note that K_D_ values from distinct studies [55, 56] might not be directly comparable. (G) Protein levels of GST-tagged Cln2 fusions expressed from promoters of different strengths (P_GAL1_, P_GALS_, P_GALL_). Expression was induced with galactose for 105 min. G6PDH served as a loading control.

**Figure S2.**
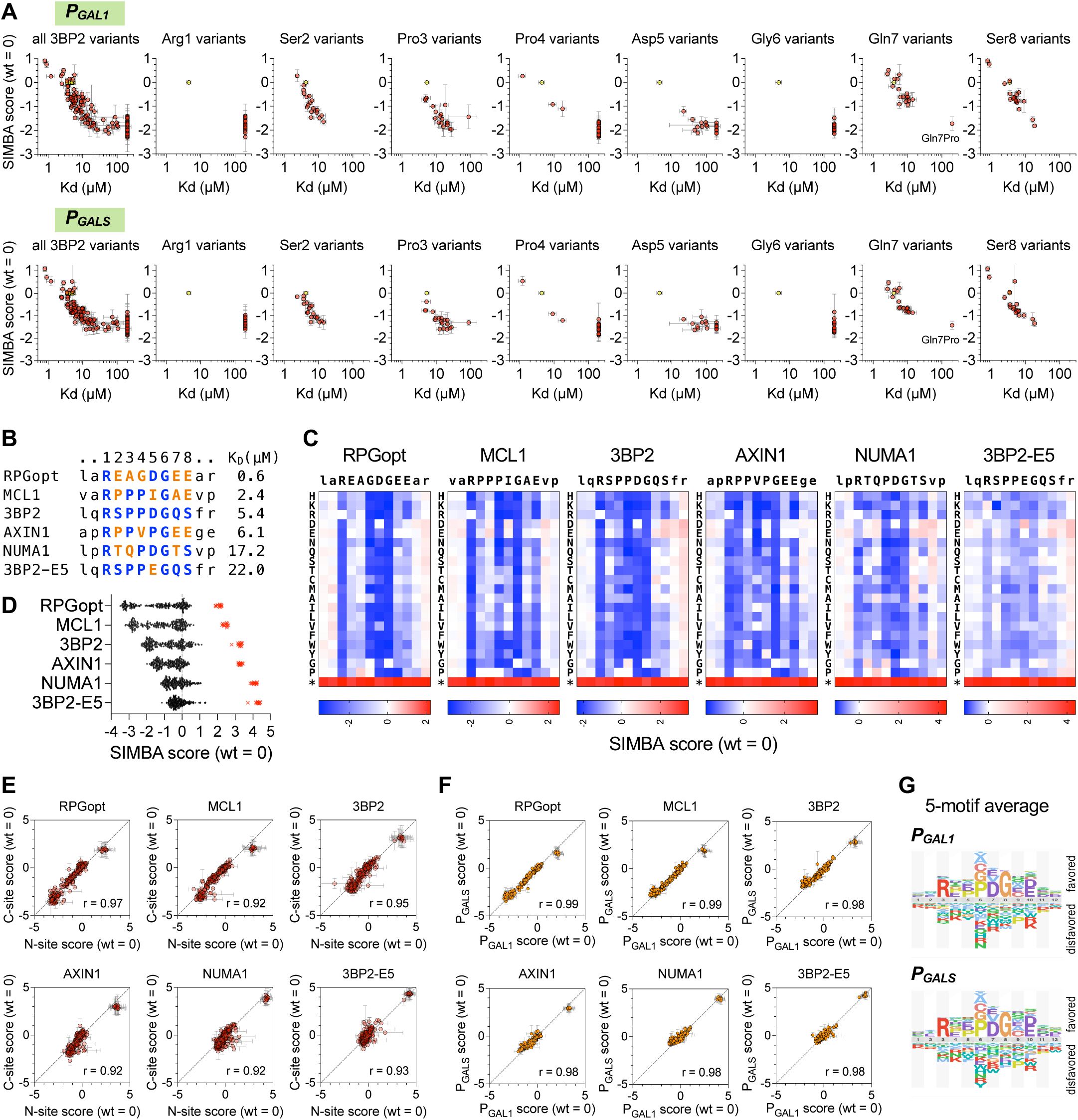
SIMBA interrogation of multiple TNKS2^ARC4^-binding peptides. (A) SIMBA scores vs. K_D_ for all 3BP2 variants as well as variants at each of 8 individual positions, plotted as in Figure 3C, with the TNKS2^ARC4^-Cln2 fusion driven by P_GAL1_ or P_GALS_. All SIMBA scores are mean ± SEM (n = 4). (B) Sequence and K_D_ of each motif subjected to DMS and SIMBA. Residues in the 8 core positions are colored where identical to (blue) or different from (orange) the 3BP2 sequence. (C) Heatmaps of DMS results from six parent peptides, plotted as in Figure 3A. (D) Distribution of SIMBA scores for all missense variants (black circles) and the average STOP codon (red X symbols) at the 12 positions in each of the 6 parent motifs. (E) Correlation of SIMBA scores for variants inserted a the N-site vs. C-site locations, plotted as in Figure 3G, for six parent peptides. (F) Correlation of SIMBA scores when the TNKS2^ARC4^-Cln2 fusion was expressed from P_GAL1_ vs. P_GALS_, plotted as in Figure 3F, for six parent peptides. (G) Sequence preference logos, averaged from 5 motifs (excluding 3BP2-E5), comparing when the TNKS2^ARC4^-Cln2 fusion was expressed from P_GAL1_ vs. P_GALS_.

**Figure S3.**
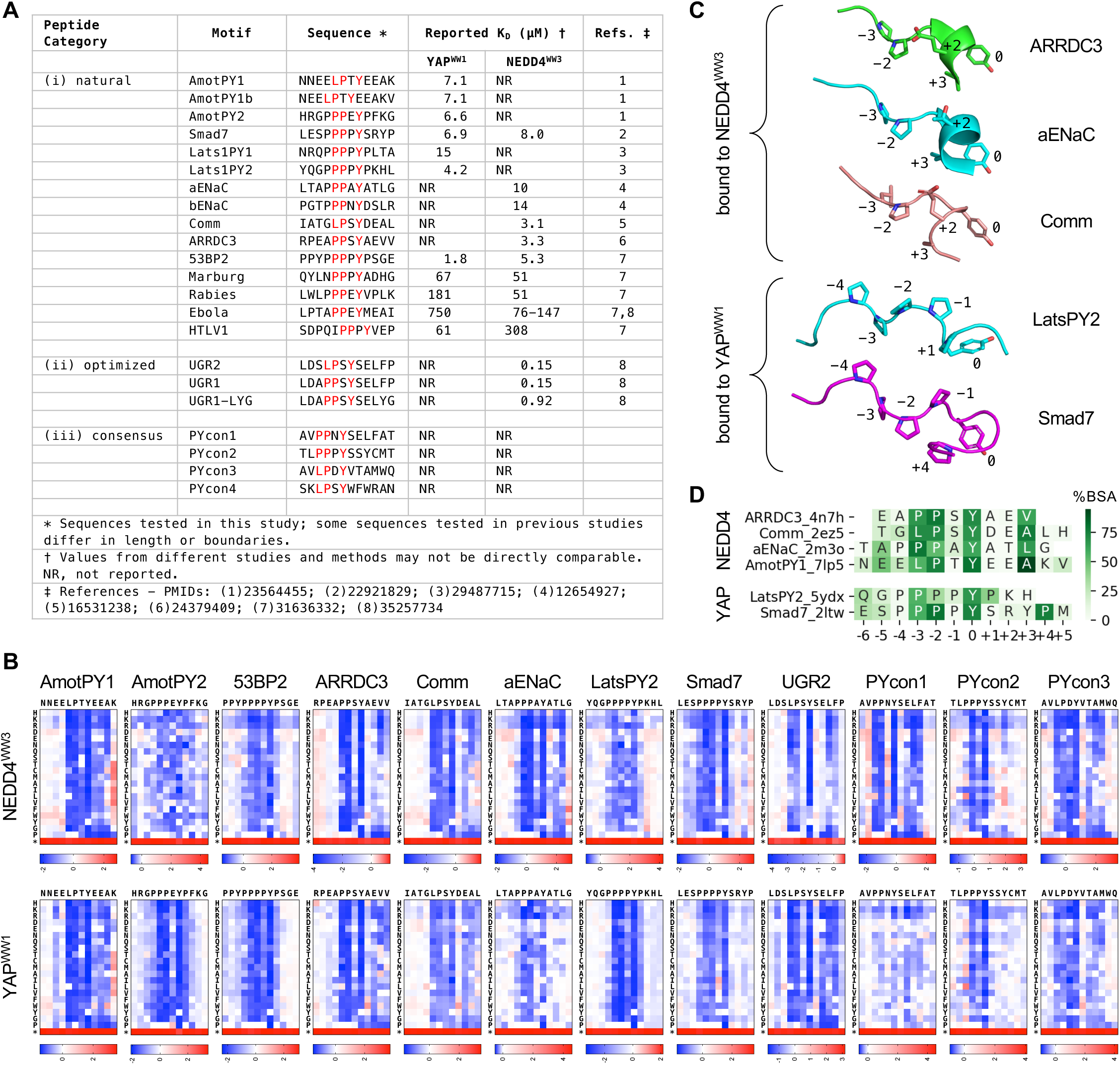
DMS data and characteristics of WW domain-binding peptides. (A) WW domain peptides used in this study, and their reported binding affinities. The consensus peptides (PYcon1-4) were designed based on “multiple PWM” sequences from Figure S1B of Ref [74], which were proposed as improved predictors of binding specificity compared to single PWMs. (B) Heatmaps of DMS results for 12 peptides combined with 2 WW domains. Data are the mean of 3 independent replicate experiments. (C) Conformations of WW domain bound peptides in published structures. The orientation is comparable to that in Figure 6G. PDB IDs: 4n7h, 2m3o, 2ez5, 5ydx, 2ltw. (D) Plot of buried surface area (BSA) for residues in WW-bound peptides, expressed as a percentage of accessible surface area in the unbound peptide. Data are from 4 NEDD4^WW3^ complexes (all type 1) and 2 YAP^WW1^ complexes (type 2). Peptide names include PDB IDs.

### Supporting Tables

**Table S1.**
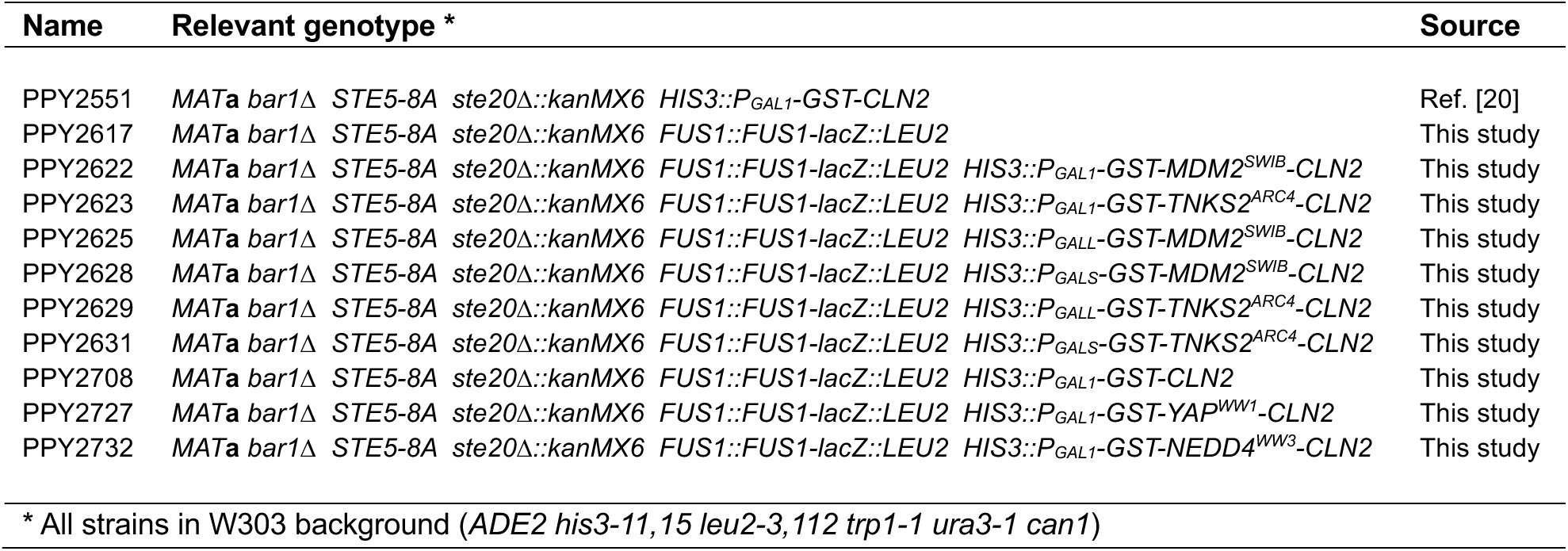
Yeast strains used in this study.

**Table S2.**
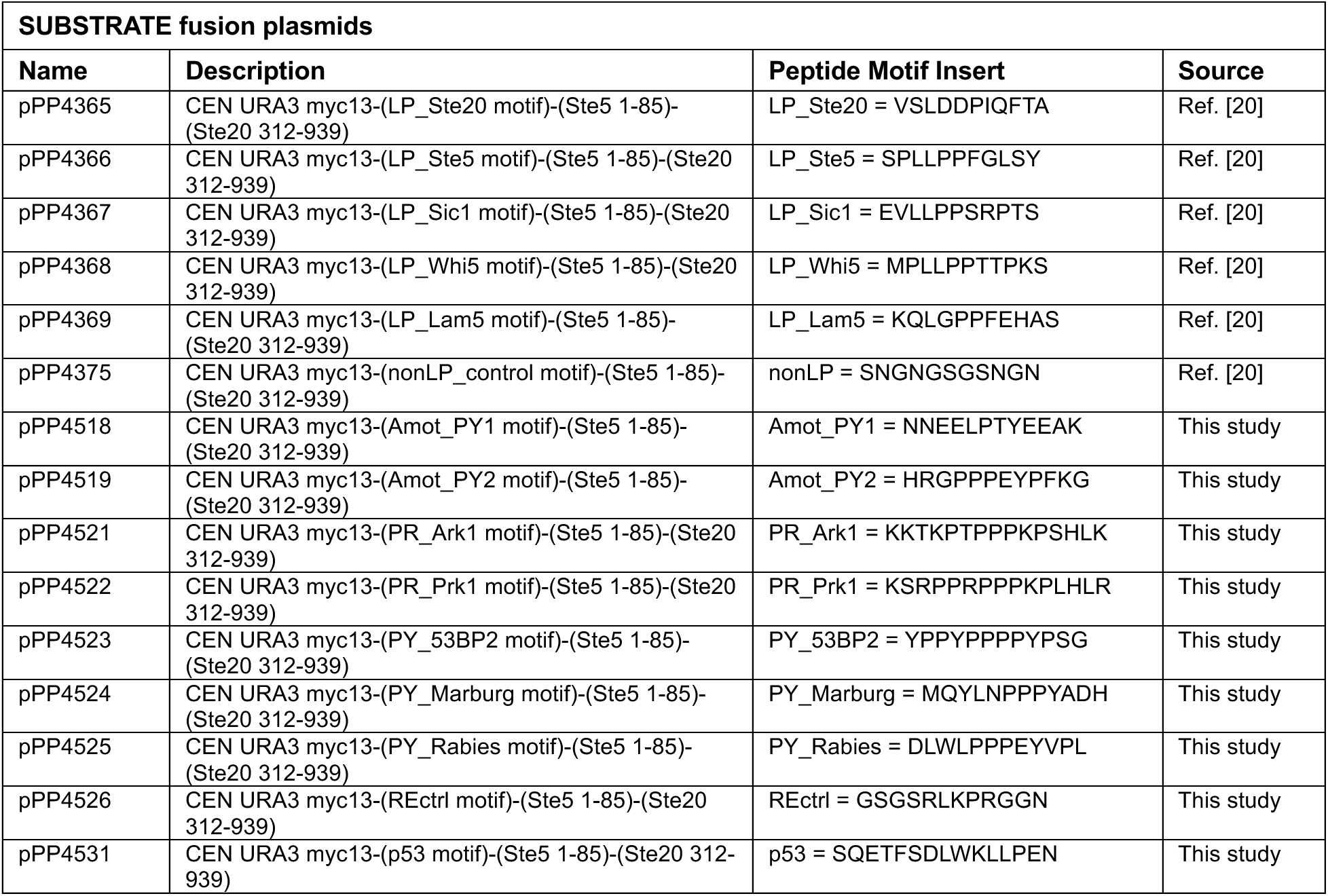
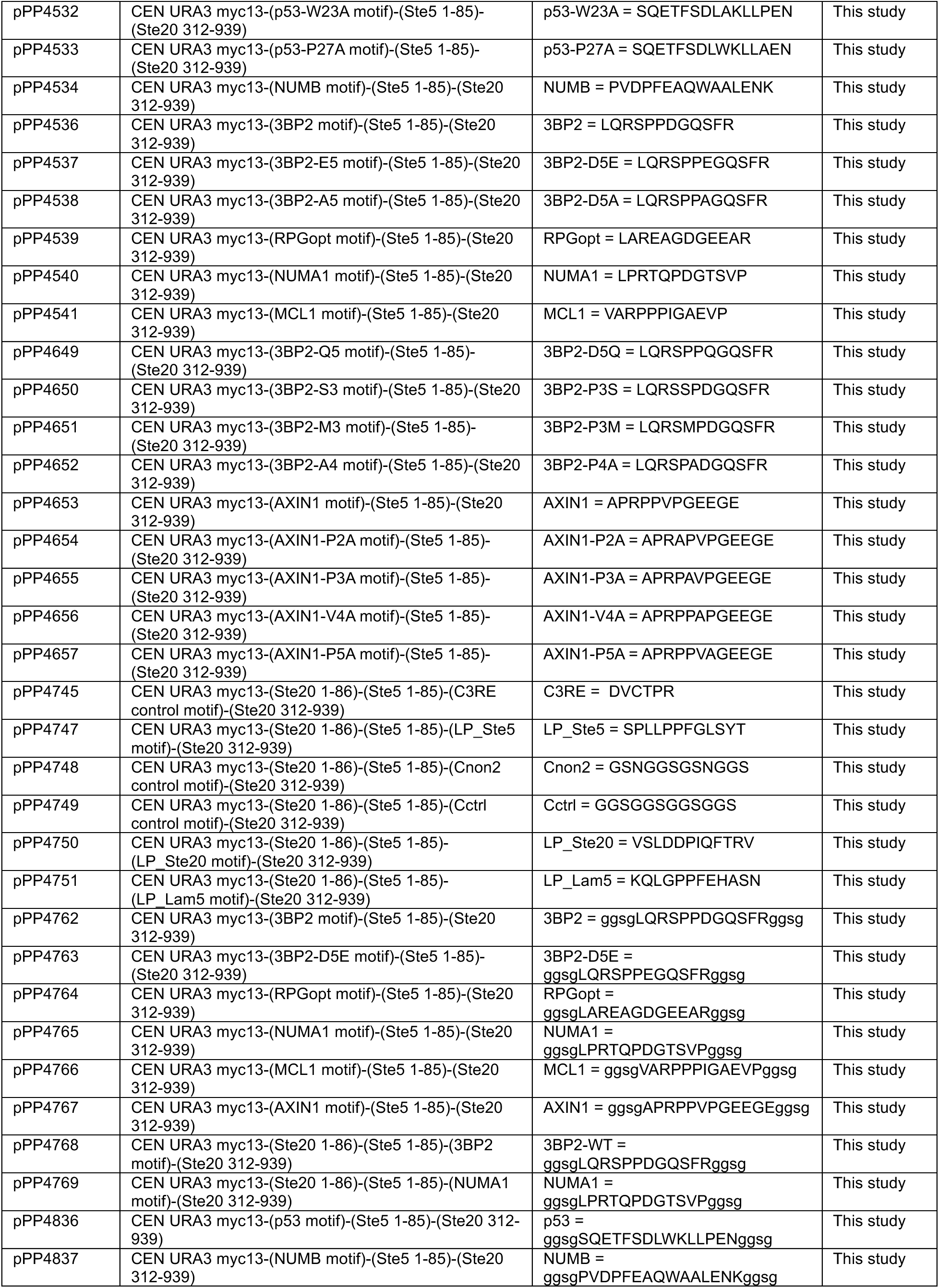
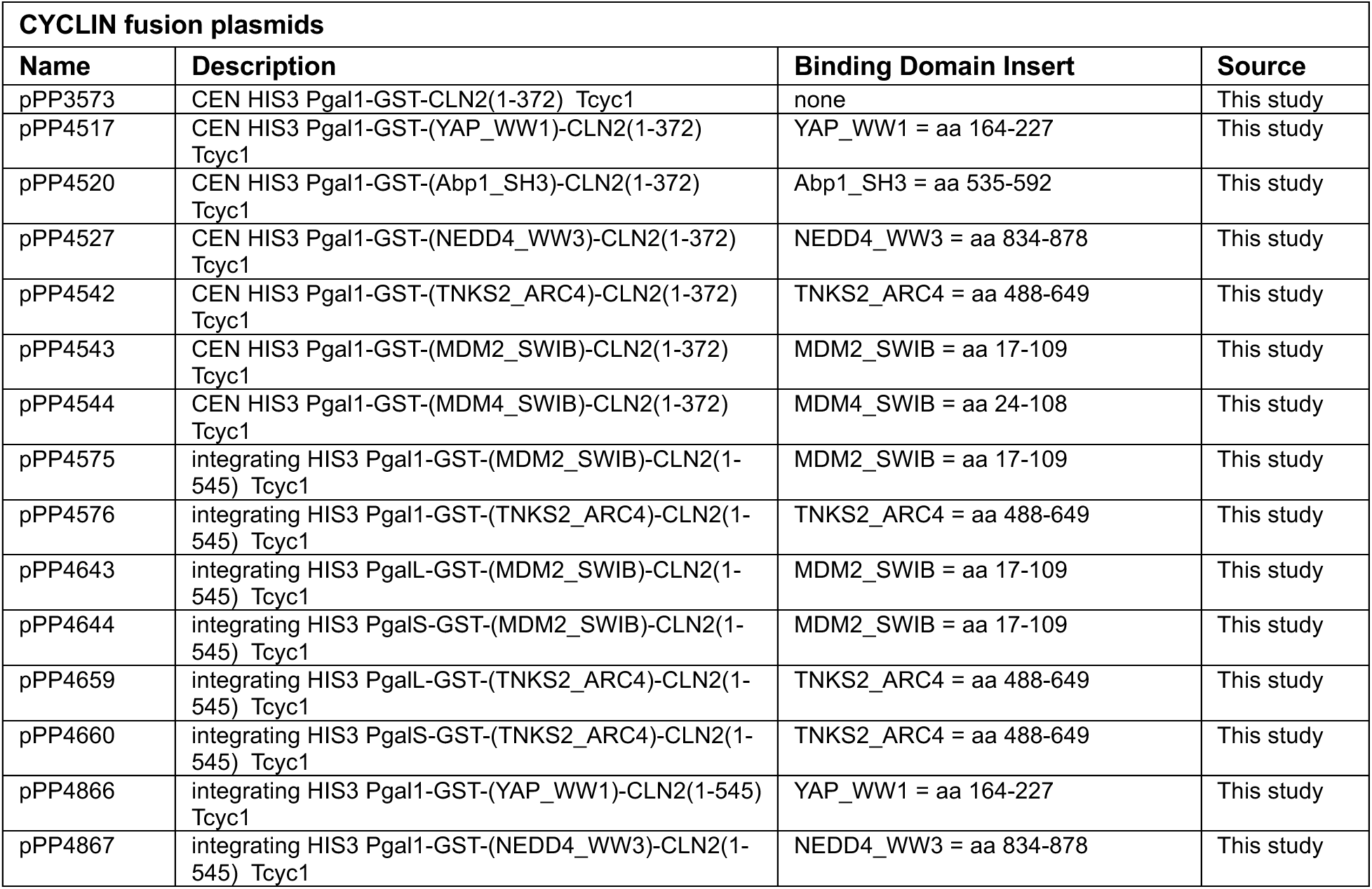
Plasmids used in this study.

